# Post-glacial recolonization and multiple scales of secondary contact contribute to contemporary Atlantic salmon (*Salmo salar*) genomic variation in North America

**DOI:** 10.1101/2023.08.28.555076

**Authors:** Cameron M. Nugent, Tony Kess, Barbara L. Langille, Samantha V. Beck, Steven Duffy, Amber Messmer, Nicole Smith, Sarah J. Lehnert, Brendan F. Wringe, Matthew Kent, Paul Bentzen, Ian R. Bradbury

## Abstract

**Aim:** In northern environments, periods of isolation during Pleistocene glaciations and subsequent recolonization and secondary contact have had a significant influence on contemporary diversity of many species. The recent advent of high-resolution genomic analyses allows unprecedented power to resolve genomic signatures of such events in northern species. Here, we provide the highest resolution genomic characterization of Atlantic salmon in North America to infer glacial refugia and the geographic scales of postglacial secondary contact.

**Location:** North America.

**Taxon:** Atlantic salmon, *Salmo salar*.

**Methods:** Samples were collected for 5455 individuals from 148 populations encompassing the majority of Atlantic salmon’s native range in North America, from Labrador to Maine. Individuals were genotyped using a 220K SNP array aligned to the Atlantic salmon (*Salmo salar*) genome. Spatial genetic structure (PCA, k-means clustering, admixture) was evaluated in conjunction with genomic comparisons of identified lineages to infer the refugia during the last glacial maximum and regions of secondary contact following recolonization.

**Results:** Spatial genomic analyses identified three phylogeographic groups, consistent with the northward recolonization from two southern glacial refugia in North America (a western Maritime lineage and an eastern Newfoundland and Labrador lineage), with subsequent differentiation of the eastern lineage into two separate groups. Secondary contact among these North American groups was present within the northern Gulf of St. Lawrence and evidence of trans-Atlantic secondary contact was detected within the eastern Newfoundland and Labrador lineage. Comparison of groups from insular Newfoundland with those from mainland Labrador suggests genomic regions displaying high differentiation were characterized by elevated European admixture, suggesting a possible role of European secondary contact in population divergence.

**Main Conclusions:** These findings present the first evidence suggesting that genomic diversity in extant North American Atlantic salmon populations has resulted from allopatric isolation in two glacial refugia followed by both regional and trans-Atlantic recolonization and secondary contact and demonstrate the power of genomic tools to resolve historical drivers of diversity in wild populations.

## INTRODUCTION

Numerous processes can shape patterns of genomic diversity within species, including: changes in species distributions (*e.g.*, Antão *et al*. 2022), secondary contact among lineages (e.g., Bernatchez 1997; Moore *et al*. 2015), and both historical and contemporary environmental adaptation (Sylvester *et al*. 2018; Watson *et al*. 2021; Pratt *et al*. 2022; Brauer *et al*. 2023). Across the northern hemisphere, Pleistocene glaciations have resulted in periods of isolation and allopatric divergence of both terrestrial and marine populations, with genomic signatures of these events persisting even following secondary contact and subsequent admixture (Pinceel *et al*. 2005; Gompert *et al*. 2010; Bierne *et al*. 2011; Bernatchez & Wilson 1998; Brunner *et al*. 2001; Rougemont & Bernatchez 2018; Saarman *et al*. 2019; Ito *et al*. 2020). As these longstanding periods of isolation have been shown to be associated with both neutral and adaptive divergence among lineages (Duranton *et al*. 2018; Stanley *et al*. 2018; Knutsen *et al*. 2022; Pratt *et al*. 2022; Nugent *et al*. 2023), they may shape contemporary diversity and responses to stressors such as climate change (e.g., Rendon-Anaya *et al*. 2021; Luqman *et al*. 2023). Ultimately, resolving the scale and drivers of diversity present in wild populations (such as colonization history and secondary contacts among glacial lineages) remains an ongoing challenge in many species and a lack of clear information can hinder species conservation and biodiversity management (e.g., Kardos *et al*. 2021).

Across the North Atlantic Ocean, genetic and genomic studies of population structure in marine and anadromous species over the last few decades have revealed large-scale patterns consistent with isolation in multiple refugia during the last Pleistocene glaciation (*e.g.*, Maggs *et al*. 2008; Provan 2013; Bringloe *et al*. 2022). At broad scales, evidence for trans-Atlantic divergence in many species examined from macroalgae to marine fishes (Maggs *et al*. 2008; Li *et al*. 2015, Lehnert *et al*. 2019) provide evidence that species persisted on both sides of the North Atlantic during the last glacial period. These studies support significant roles of glacial isolation in driving contemporary diversity, both at trans-Atlantic and local scales, across a diverse suite of aquatic taxa and suggest recolonization from southern refugia in both the eastern and western Atlantic (Wares 2001; Maggs *et al*. 2008; Bernatchez 1997) as well as the possibility of high latitude periglacial refugia (Maggs *et al*. 2008; Bringloe *et al*. 2022). Genomic evaluations now permit the impacts of isolation and post-glacial recolonization to be quantified genome-wide and have revealed signatures of glacial isolation as well as evidence of post glacial secondary contact and introgression among lineages (*e.g.*, Souissi *et al*. 2018; Lehnert *et al*. 2019; Salisbury *et al*. 2023). These studies support the hypothesis that Pleistocene glaciations have been a significant determinant of contemporary diversity in northern marine species, which is likely to influence population and species response to climate change in the coming decades (e.g., Luqman *et al*. 2023).

Atlantic salmon (*Salmo salar*) is an anadromous salmonid species distributed throughout the North Atlantic that is of significant conservation concern due to drastic demographic declines in recent decades across much of its natural range (Lehnert *et al*. 2019). Population structure of Atlantic salmon has repeatedly been shown to be hierarchical in nature, with large trans-Atlantic differences contrasting regional and river specific genomic structure resulting from the species’ strong homing behaviour (Dodson *et al*. 1998; Fraser *et al*. 2011; Moore *et al*. 2014; Wellband *et al*. 2018). Existing evidence suggests extensive genome-wide differences between European and North American populations despite evidence of secondary contact in northern portions of the North American and European ranges (Makhrov *et al*. 2005; Bradbury *et al*. 2015; Rougemont & Bernatchez 2018; Lehnert *et al*. 2019). Within North America, previous microsatellite and small SNP array-based studies suggest extensive regional and river scale divergence (e.g., Moore *et al*. 2014; Bradbury *et al*. 2021) within two large scale North American groups (Rougemont & Bernatchez 2018). Post-glacial colonization history has been examined in other geographic regions; for example, analysis of Atlantic salmon throughout the Baltic Sea has inferred postglacial recolonization by two phylogeographic lineages (Koljonen *et al*. 1999). However, the genomic consequences of glacial isolation and potential secondary contact have not been quantified using high density genomic data in the Northwest Atlantic beyond examination of trans-Atlantic divergence (see Lehnert *et al*. 2019, 2020).

The goals of this study were to provide a high-level characterization of Atlantic salmon genomic variation and population structure in the Northwest Atlantic, based on the largest set of individuals examined on a genome-wide scale to date. To accomplish this, we leveraged a previously published data set of 5455 individuals from 148 sampling locations in northeastern North America (Lehnert *et al*. 2020; Bradbury *et al*. 2022; Nugent *et al*. 2023), where individuals were genotyped using a high-density (220K) SNP array that provides extensive coverage of all chromosomes (Barson *et al*. 2015; Lehnert *et al*. 2019). We used these data to infer the demographic history of the species in the northwest Atlantic region within the context of the most recent deglaciation of the region and historical European secondary contact. To achieve this, we: *i*) characterized genomic diversity and population structure across the sampled range of locations throughout northeastern North America, *ii*) analyzed the phylogeographic relationships of populations, *iii*) searched for evidence of admixture between major genetic groups, and *iv*) used these data to make inferences about the demographic history of Atlantic salmon in the study region. This research builds upon previous Atlantic salmon population genomics studies and provides novel insights into the high-level population structure of the species and how it has been influenced by past climate change.

## METHODS

### Samples and genetic information

A series of 5455 Atlantic salmon were collected from 148 rivers throughout northeastern North America (Figure 1; Supplementary Figure S1). The number of samples from each location ranged from 9 to 209, with a mean of 41. These samples were genotyped using a 220K bi-allelic Affymetrix Axiom SNP array developed for use in Atlantic salmon (Barson *et al*. 2015). The samples were used in previously published sources and the methods for DNA extraction, genotyping, and bioinformatics pipelines for SNP data quality control can be found therein (Lehnert *et al*. 2020; Bradbury *et al*. 2022; Nugent *et al*. 2023). Briefly, these processing steps involved filtering genotype data for high quality SNPs based on their clustering patterns and retaining only SNPs with “polymorphic high” assignments and call rates >0.99. Using *Plink* (version 1.9; Chang *et al*. 2015), SNPs were then filtered for a minimum minor allele frequency (MAF) cutoff of 0.01 across loci and SNPs were pruned based on linkage disequilibrium using the parameters:*--indep-pairwise 50 5 0.5*.

**Figure 1.**
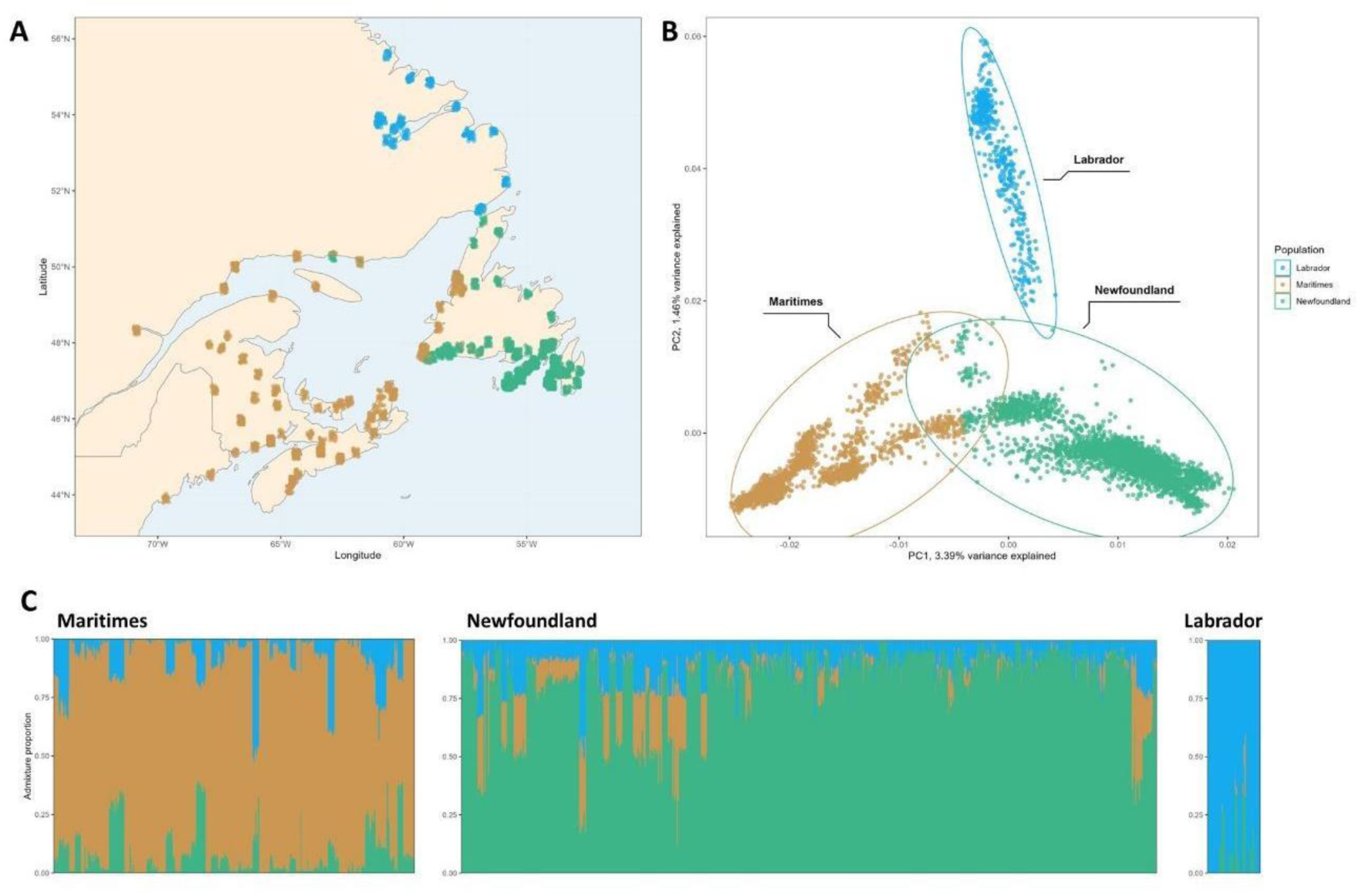
Broad-scale genetic structure North American Atlantic salmon populations inferred by *k*-means clustering and admixture analysis of 5455 individuals based on 65576 SNPs. A) Map of sampling locations showing the predominant genetic cluster at each location, with each point representing a sample. B) Scatter plot of Principal Components (PCs) of genetic variation; colours correspond to the predominant cluster to which individuals were assigned. C) Per-individual admixture proportions.

The location of the 220K array SNPs within the latest version of the Atlantic salmon reference genome (version: Ssal_v3.1, RefSeq Number: GCF_905237065.1) were determined. To do this, known SNP positions and 100bp sequence flanks were excised from the previous version of the Atlantic salmon genome (version: ICSASG_v2, RefSeq Number: GCF_000233375.1). For SNPs with no known location in the previous genome assembly, the Affymetrix SNP primer sequences were utilized. The program *bwa* (version 0.7.17, Li & Durbin 2009) was used to conduct a Burrows-Wheeler alignment of these sequences against the newest Atlantic salmon reference genome. The resulting alignment information then processed using a custom Python program, *snp-placer* (https://github.com/Cnuge/snp-placer), to determine the exact base pair location of the array SNPs within the newest Atlantic salmon reference genome (version: Ssal_v3.1).

### Characterization of genomic variation via principal component analysis

To quantify population structure, a principal component analysis (PCA) of the filtered genotype data for the 5455 Atlantic salmon samples (65576 SNPs post filtering) was conducted using the R package *PCAdapt* (Luu *et al*. 2017) with the parameters *k* = 2 and *min.maf* = 0.01; the resulting principal components (PCs) were visualized in R using the package *ggplot2* (Wickham., Chang, & Wickham, 2016). To identify the regions of the genome most significantly contributing to the genomic differences described by the PCA, the per marker *p* values and PC loading scores were visualized using Manhattan plots and inspected for evidence of peaks within the genome.

To quantify putative population structure, *k*-means clustering of values on the first and second PC axes was iteratively conducted for values of *k* ranging from 1 to 10; for each value the within cluster sum of squares (*wss*) was calculated. A scree plot was produced and used to determine the optimal *k* value for the data through identification of the inflection point in the *wss* trendline. Final *k*-means clustering using the optimal value of *k* was conducted and individuals were labelled as members of putative groups based on their cluster assignments. To look for evidence of hierarchical structuring, additional PCA analyses were run for individuals within the identified groups, using the previously described methods.

### Characterizing admixture of groups

Admixture analysis of the complete set of samples was conducted to further explore population structure, resolve the boundaries between the identified population clusters, and search for evidence of individuals of mixed genetic origin. The program *Admixture* (*version 1.3.0*; Alexander *et al*. 2009) was run using the best value of *k* as determined through *k*-means clustering analysis. The resulting per individual Q values were imported into R and visualized using *ggplot2* (Wickham *et al*. 2016). The identified clusters from the admixture analysis were mapped to the three genetic groups from the previous *k*-means clustering and the data were inspected to search for evidence of admixture in individuals and patterns relating to their geographic origin. High admixture individuals, representing likely ongoing hybridization, were arbitrarily defined as those where the Q values for their primary population assignment was less than two thirds of the entire genome. The number and frequency of high admixture individuals per location was calculated and their geographic distribution was visualized. We also searched for evidence of directional bias in admixture between groups through pairwise inspection of the correlations of admixture proportions between the groups.

### Assessing genetic differences with F_ST_

The genomic basis of differentiation among identified genetic groups was examined using pairwise comparisons of locus-specific *F*_ST_. For each pairwise comparison, *Plink* was used to subset the individuals from the complete data set and the *--fst* flag was used to calculate the per marker *F*_ST_ values. The outputs were read into R, visualized as Manhattan plots using *ggman* (https://github.com/drveera/ggman), and the genome-wide mean, minimum, and maximum *F*_ST_ values were calculated. The number of outlier loci (defined as those exhibiting *F*_ST_ greater than three standard deviations from the mean for a given comparison) were quantified and the distribution of these loci throughout the genome was determined through inspection of Manhattan plots and assessment of the *Plink* output files.

### Linkage Disequilibrium

We searched for evidence of reduced recombination throughout the genome by conducting windowed linkage disequilibrium (LD) analysis, for both the complete set of individuals and separately for each of the identified clusters. To eliminate bias, the LD analysis utilized the complete set of markers passing quality and minor allele frequency filtering (no pruning for LD was applied). For each group, *Plink* was used to calculate LD for 1Mb windows using the flags: *--ldwindow-kb 1000 –r*. A custom Python script (Supplementary File 1 of Nugent *et al*. 2023) was used to summarize the windowed LD values on a 1Mb basis, calculating the mean, median, min, and max *r^2^* as well as the sample size. The summarized outputs were imported into R and visualized as Manhattan plots. The *Plink* output files were also read in and *r^2^* values were visualized as histograms to search for evidence of outliers in the *r^2^* value distributions. Regions of high LD were defined as those with median *r^2^* values greater than 3 standard deviations from the given group’s genome-wide median score.

### Signals of ancient European admixture

We searched for evidence of ancient (*i.e.* non-anthropogenic) European admixture and its potential influence on population structure across the sampled North American range. To achieve this, we compared the per individual admixture proportions for each North American population to per individual estimates of European admixture proportions that were obtained from previous quantification in Nugent *et al*. (2023). To assess the magnitude and geographic distribution of European admixture, for each North American lineage the correlation of per individual admixture proportions for the given group with European admixture proportions were quantified following Bradbury *et al*. (2022) and Nugent *et al*. (2023).

Based on results of preliminary assessment of European admixture influences (see Results), the role of European admixture as a source of differentiation between the Labrador (LAB) and Newfoundland (NFL) phylogeographic groups was further examined. A series of admixture analyses were conducted to examine the estimated admixture of individuals from the LAB and NFL phylogeographic groups in conjunction with European samples. First, to confirm that the NFL and LAB groups were more similar to one another than to the European individuals (as would be expected based on previous analyses presented in: Lehnert *et al*. 2019, Bradbury *et al*. 2022, and Nugent *et al*. 2023), an admixture run for 431 LAB, 3244 NFL, and 804 wild Norwegian samples using all polymorphic markers (data obtained from Nugent *et al*. 2023) was conducted with the parameter *k* = 2. Admixture results using Norwegian salmon have previously been shown to demonstrate the same signal as using multiple group of European fish (Nugent *et al*. 2023). Second, to test if regions of genomic differentiation between the NFL and LAB populations displayed elevated European ancestry, the NFL, LAB, and European admixture analysis was rerun using only the 1648 *F*_ST_ outlier SNPs identified in comparison of the NFL and LAB groups. Finally, to check that the *F*_ST_ outlier loci effectively separated the LAB and NFL groups, admixture was rerun using *k* = 2, only the samples from the NFL and LAB groups, and the 1648 *F*_ST_ outlier loci. As a check to rule out assignment error, all analyses were repeated using k = 3 to ensure the three groups (NFL, LAB, and European individuals) were identified under these conditions.

To search for evidence of the European chromosome 1/23 translocation (that has previously been associated with trans-Atlantic introgression in these geographic regions, see: Lehnert *et al*. 2019; Watson *et al*. 2022) contributing to differentiation of LAB and NFL, we calculated the per chromosome frequency of *F*_ST_ outlier loci and conducted a chi-squared test to see if there was evidence of a significant difference in per-chromosome *F*_ST_ outlier rate.

Lastly, a constrained reanalysis of the genomic structure was conducted to provide an alternative test of the extent to which loci associated with European introgression were contributing to the observed structure of the described populations. Genotypes of the complete set of the 5455 North American individuals and the 804 wild Norwegian samples were analyzed with *PCAdapt* (Luu *et al*. 2017) using *k =* 2. The *p* values of loci were obtained, giving a metric of their association with the genetic separation of North American and European salmon (*e.g.* Figure 1C. of Nugent *et al*. 2023). To see if the removal of loci associated with European introgression altered the observed structure of North American Atlantic salmon, PCA of the 5455 North American samples was then repeated with the 25% of marker (n=16394) showing the highest *p* values for association with North American – European genetic structure removed.

### Reconstruction of evolutionary and demographic history

The phylogenetic relationships and evolutionary history of the sampling locations were inferred using *treemix* (version 1.13; Pickrell & Pritchard 2012). A set of 10000 SNPs were randomly selected from the complete set of LD filtered markers and a *treemix* input file was constructed, with the marker order based on the location of the SNPs in the Ssal_v3.1 Atlantic salmon reference genome; the per-location maximum likelihood tree was then constructed.

We calculated and visualized per-location average genome wide heterozygosity in order to assess the intra-population distributions of genetic diversity and look for signatures of colonization trajectories (*e.g.* test the hypothesis of lower heterozygosity at more northern latitudes). For each sampling location, *Plink* was used to subset relevant individuals and conduct per locus assessment of heterozygosity (using the *--hardy* option). In R, the data were then summarized to calculate the mean observed genome wide heterozygosity. The process was repeated on a per population basis, using all the individuals that were assigned to a given population in the *k*-means clustering analysis.

To examine recent changes in demography and compare observed patterns across the identified phylogeographic groups, the effective population size (N_e_) of the identified phylogeographic groups was estimated using GoNE (Genetic Optimization for N_e_ Estimation; Santiago *et al*. 2020). First, *Plink* was used to subset the individuals of the relevant clusters from the complete data set and produce a PED and MAP file for each of the identified populations. For each population, the data were then further subset to limit N_e_ estimation to the regions where the majority of individuals did not display high within-North America admixture (Table S1) as admixture can introduce artefacts into N_e_ estimates (Santiago *et al*. 2020). The NFL population was then further randomly down sampled to 1700 individuals in order to fit the maximum population size that GoNE can accommodate (Santiago *et al*. 2020). The inputs were then analyzed with GoNE using default parameters and the output files were inspected and visualized in R using *ggplot2* (Wickham *et al*. 2016).

### Gene ontology

The annotation of the Atlantic salmon genome (version Ssal_v3.1, RefSeq Number: GCF_905237065.1) was queried to search for evidence of genes associated with loci or regions of interest identified in previous analyses. The location of SNPs identified as significantly associated with the PCs from the PCA of genomic variation and SNPs with an *F*_ST_ greater than three standard deviations from the mean for any of the pairwise combinations of differentiation analyses were assessed to identify SNPs of interest that were positioned between the start and end point of annotated genes. For regions of the genome defined as having high LD, the high LD windows were compared to the start and end points of known genes to identify any genes that overlapped the windows of interest (possessing a start or end point within the given window of the genome). The three sets of results were collated to search for evidence of genes displaying evidence of association with more than one of the statistical tests.

The collated significant SNPs were then subjected to Gene Ontology (GO) analysis in order to elucidate any enriched gene pathways. The list of Entrez GeneIDs was loaded into ShinyGo (version 0.76.3; http://bioinformatics.sdstate.edu/go/) and the GO analysis was then conducted using the pathway database option “all available gene sets” and default parameters otherwise. Enrichment information and data visualizations produced by ShinyGo were then examined to determine significantly associated gene pathways.

## RESULTS

### Characterization of Genomic Variation

The PCA of filtered genotype information for the 5455 Atlantic salmon samples revealed evidence of geographically correlated structure; the first principal component (PC) axis explained 3.39% of genomic variance and the second PC explained 1.46% of variation (Figure 1). There were 485 SNPs with Bonferroni corrected *p* values < 0.05, indicating significant association with the PCs. Inspection of *p* values characterizing each marker’s association with the PC axes revealed several regions of the genome that appeared to be significantly associated with the observed genetic structure (Figure 1; Figure 4). The *k-*means clustering of PC1 and PC2 values resolved individuals into three clusters that had distinct geographic distributions, with some overlap in both the geographic and genetic boundaries (Figure 1). The three clusters were named as putative phylogeographic groups based on their current geographic distributions: Labrador (LAB), Maritimes (MAR), and Newfoundland (NFL). Subsequent intra-cluster PCAs for each of the three phylogeographic groups showed evidence of hierarchical structuring, as would be expected of salmon populations (Supplementary Figure S2).

### Evidence of admixture

The admixture analysis displayed strong concordance with the *k*-means clustering analysis; all individuals had admixture values of Q > 0.5 for their identified cluster (Figure 1). There was evidence of genetic admixture, with 17.7% of individuals displaying Q values of less than 0.66 for their home cluster, thereby suggesting mixed genetic origins (Figure 2). These high admixture individuals were concentrated to specific sampling locations, with only 44/148 locations containing one or more highly admixed individual and 29/148 locations where more than half of the sampled individuals were classified as high admixture (Figure 2). These high admixture locations represented the putative geographic boundaries between the three main clusters, suggesting that there is admixture of genetic groups in putative transition zones between phylogeographic groups (Wennevik *et al*. 2019). Examination of pairwise scatter plots of admixture proportions revealed evidence of non-homogenous admixture across the phylogeographic groups (Figure 6). Directional bias was observed for some combinations: bidirectional admixture between MAR and NFL groups, strong evidence of admixture from LAB to MAR but weak evidence of the inverse, and strong evidence of admixture from NFL to LAB but weak evidence of the inverse (Figure 6).

**Figure 2.**
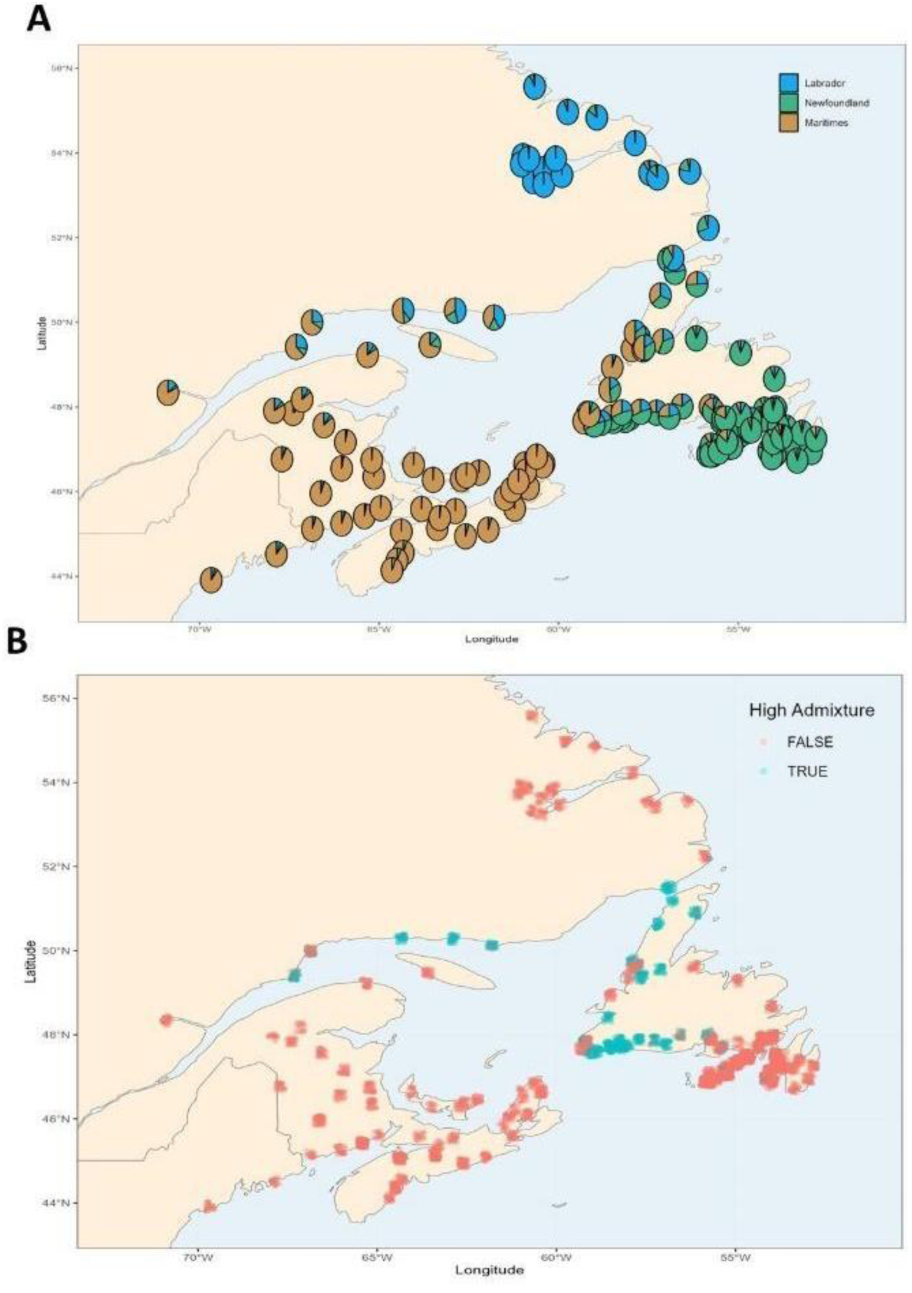
Geographic variation in genetic admixture of Atlantic salmon populations in North America. A) Map displaying per location genetic admixture values. Each pie chart represents the cumulative admixture proportions of all individuals from a sampling location. B) Per-individual admixture proportion estimates expressed relative to a binary threshold, where the location of points shows the sampling origin of each individual, Q>0.66 is considered low admixture and Q<0.66 is considered high admixture.

**Figure 3.**
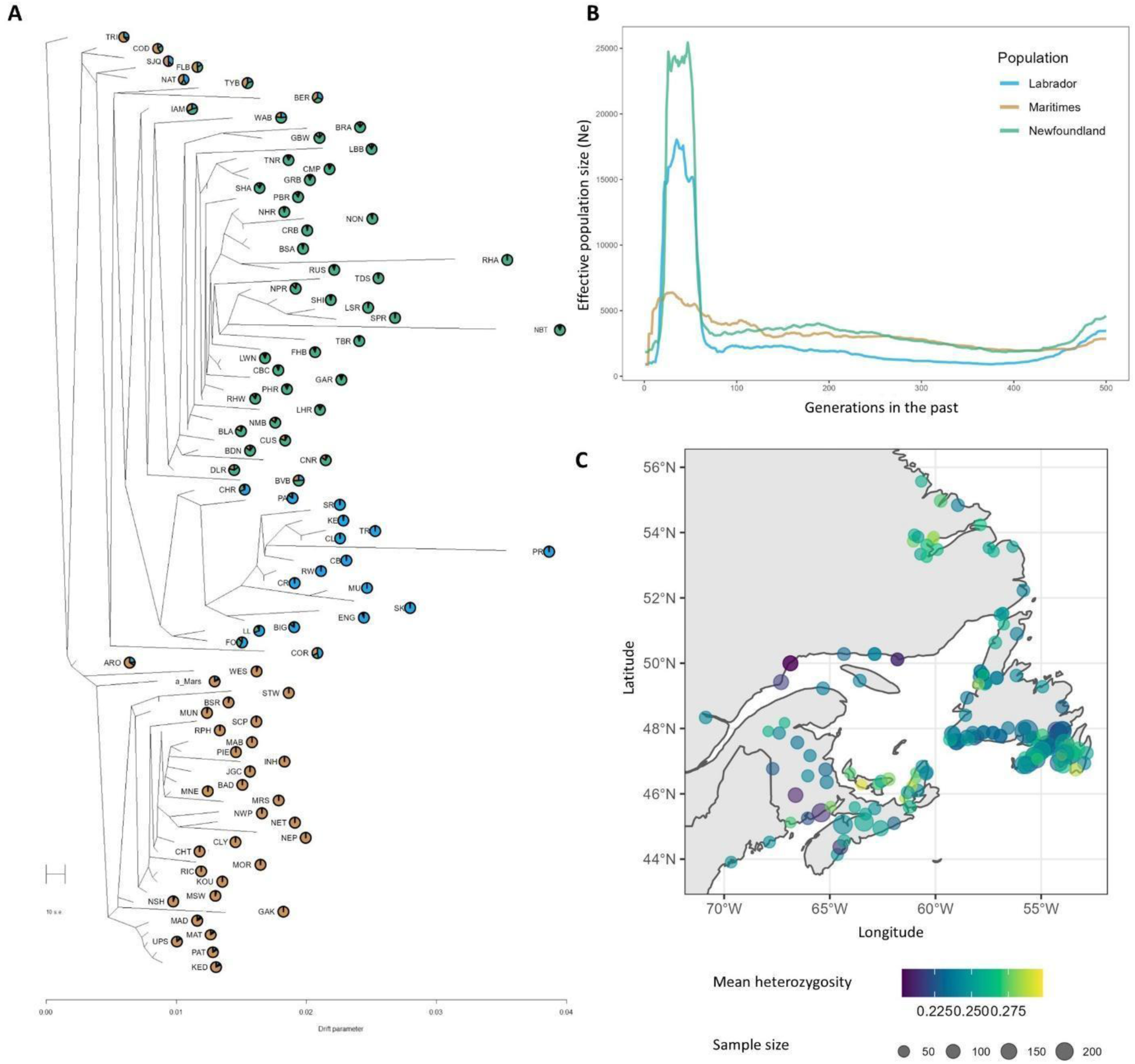
A) Phylogenetic tree of all sampling locations with > 10 individuals. The pie charts adjacent to the location labels represent the cumulative admixture proportions of all individuals from the given location. Residual values corresponding to the tree can be found in Supplementary Figure S4. B) Historical effective population size estimates produced using GoNE. C) Per-location mean heterozygosity for all sampling locations. The colour of the points indicates the genome-wide mean heterozygosity for the sampling location and the size of the points indicate the sample size for the given location.

### Assessing genomic differentiation and linkage disequilibrium

For each pairwise combination of clusters, per-locus *F*_ST_ scores were calculated to explore the genomic distribution of differentiation (Figure 4). The results revealed high levels of genome wide *F*_ST_ among the different pairs of phylogeographic groups, with all comparisons possessing mean genome wide *F*_ST_ values >0.03 and over 2000 loci exhibiting *F*_ST_ greater than three standard deviations from the comparison mean (Figure 4; Table 1). The highest levels of differentiation were observed between the LAB and MAR groups, where 6143 (9.4%) loci had *F*_ST_ values greater than 3 standard deviations from the mean, while the lowest levels of differentiation were seen between the MAR and NFL groups, where only 2097 (3.2%) loci had *F*_ST_ greater than 3 standard deviations from the mean. At regions of the genome significantly associated with observed genetic structuring there was evidence of elevated differentiation between LAB and the two other groups (Figure 4).

**Figure 4.**
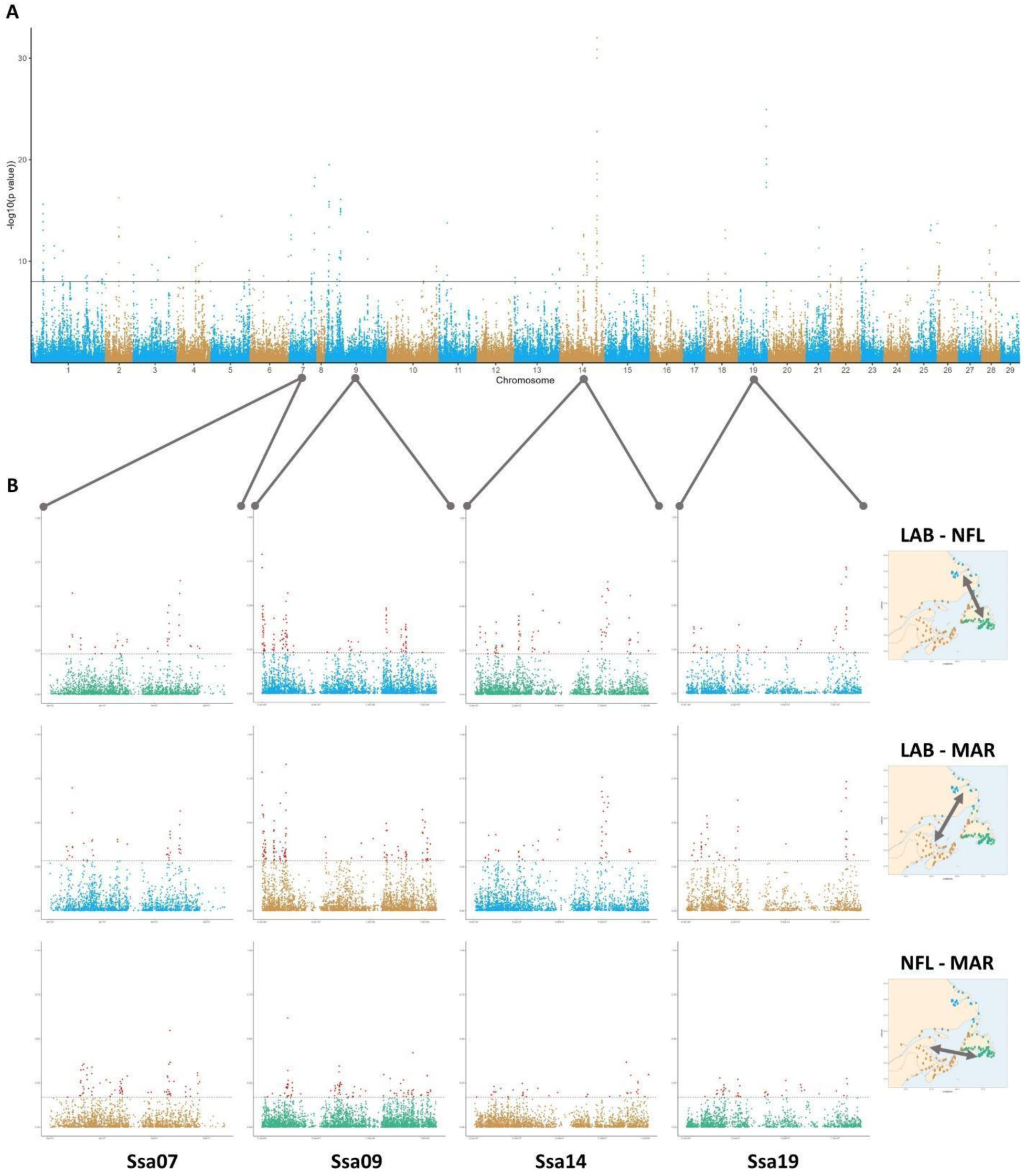
A) Manhattan plot showing the P values indicating the association of each SNP marker with the PC values of Figure 1B. B) Pairwise *F*_ST_ comparison of regions of the four chromosomes with the lowest observed P values. Horizontal lines represent a genome-wide threshold of 3 standard deviations from the mean for the given pairwise comparison, with SNPs exceeding these thresholds being coloured in red. The map to the right of each row indicates the pairwise comparison of populations represented.

**Table 1.** Summary statistics of the genomic differentiation (*F*_ST_) of all pairwise comparisons of identified Atlantic salmon phylogeographic groups. The table gives the number of loci with an *F*_ST_ value greater than 3 standard deviations (n_loci_ F_ST_ > mean +/- 3 sd), the mean genome wide F_ST_ for a given comparison (μ F_ST_) and the maximum *F*_ST_ observed for the comparison (Max F_ST_). The vertical and horizontally aligned population designations indicate the pairwise comparison of populations that corresponds to a given cell.

|  |  |  |  |
| --- | --- | --- | --- |
|  | Labrador |  |  |
| $n_{loci} F_{ST} > \text{mean} \pm 3 \text{ sd}$ | 6144 | Maritimes | |
| $\mu F_{ST}$ | 0.050 | | |
| Max $F_{ST}$ | 0.833 | | |
| $n_{loci} F_{ST} > \text{mean} \pm 3 \text{ sd}$ | 3892 | 2097 | Newfoundland |
| $\mu F_{ST}$ | 0.039 | 0.033 | |
| Max $F_{ST}$ | 0.790 | 0.618 | |

Across the complete data set and the three identified phylogeographic groups, the windowed linkage disequilibrium (LD) analysis identified 31 windows with evidence of high LD that were based on the comparison of 5 or more loci (Figure S5), which represent putative regions of reduced recombination in the genome. Of these, identified high LD windows, 19 were based on five or more comparisons of loci: 17 windows from the full data set and 2 windows from within the NL cluster (Table S2; Figure S5). Collation of results with the *F*_ST_ scores identified 15 regions of the genome from 7 unique chromosomes that displayed evidence of elevated LD as well as high levels of genome wide *F*_ST_ (Table S4.).

### Influence of ancient European introgression on population structure

Across the complete set of North American individuals, the observed proportions of European admixture were low (0.00-0.059) and appeared to follow a Poisson distribution. The comparison of European admixture proportions within the three North American lineages revealed an association with geography, with the mean European proportions varying across the phylogeographic groups: MAR = 0.006, NFL = 0.017, and LAB = 0.026 (Figure 5). Results were suggestive of historical European secondary contact along the Eastern range of sampling locations (NFL and LAB) with the entire LAB group and southeast NFL sampling locations showing evidence of low but consistent levels of ancient European admixture. There was minimal evidence of European admixture within the MAR population. The LAB population showed a strong and consistent signal of European secondary contact, with the majority of LAB-assigned individuals displaying 2-4% European ancestry. The European admixture signal within the NFL population was more varied and geographically localized, with some individuals from southeast sampling locations displaying 2-6% European ancestry (Figure 5).

**Figure 5.**
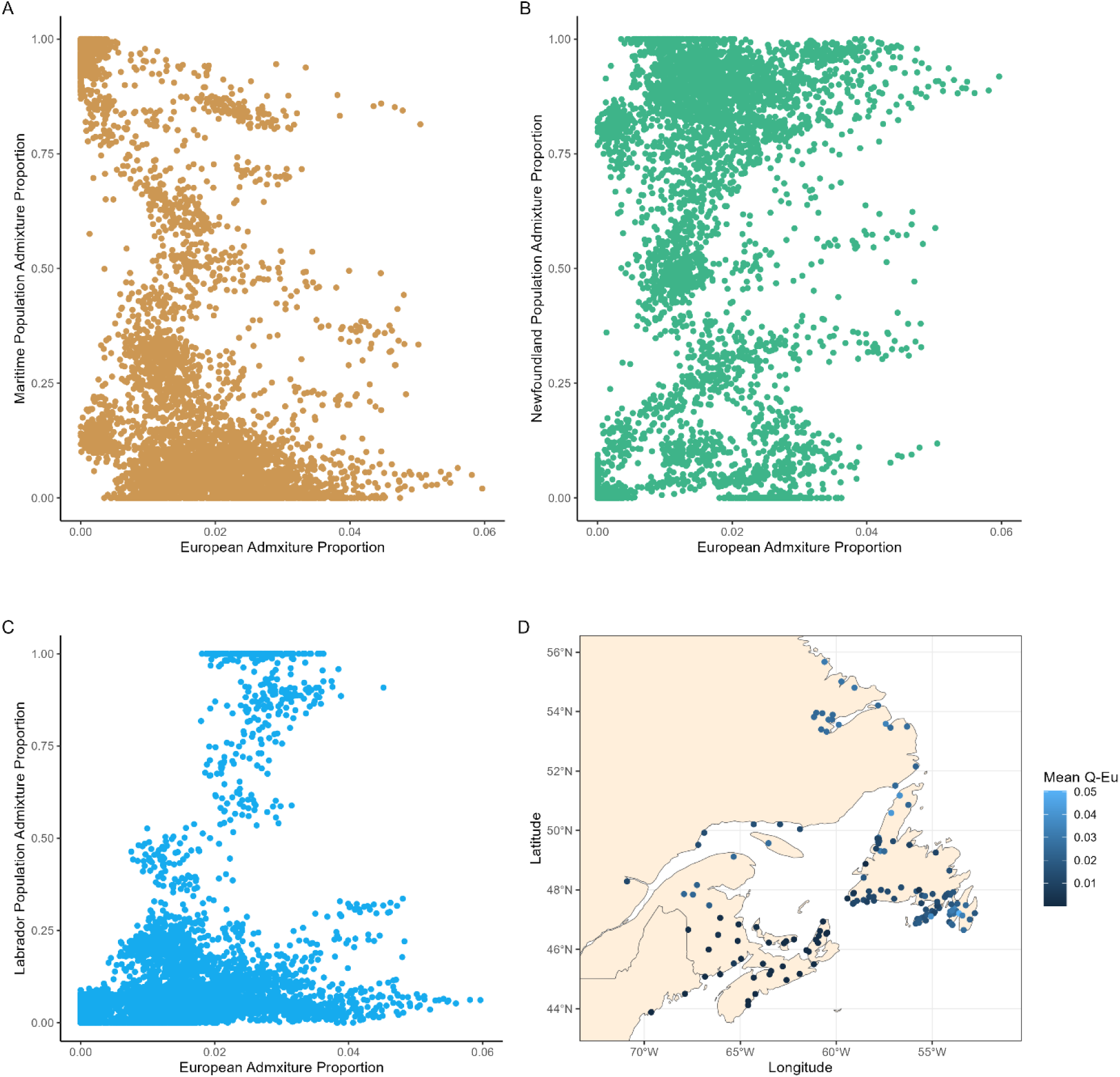
Scatter plots showing the per population admixture proportions (y) for each individual relative to their per-individual European admixture proportions (x). Plots included all of the sampled individuals (n = 5455) and their admixture proportions for the given phylogeographic group: A) MAR, B) NFL, and C) LAB. D) Map showing the sampling locations, where the colour of dots represents the per location mean European Ancestry proportion.

We explored European admixture within the NFL and LAB groups as a factor contributing to the genomic structure of these populations. First, an admixture run containing NFL, LAB, and a series of 804 Norwegian (NOR) individuals was able to replicate the European and North American genetic structure reported in Bradbury *et al*. (2022) and Nugent *et al*. (2023), with strong separation of North American and European salmon and low European admixture percentages for the LAB population (Figure 6A). Re-analysis of admixture with the same individuals and only the 1634 *F*_ST_ outlier loci from comparison of the LAB and NFL groups showed a different signal of admixture, with individuals from the LAB population being assigned much higher proportions of estimated European admixture (Figure 6B). Given the null hypothesis of no role of European ancestry in North American phylogeographic differentiation, we would expect a random distribution of admixture regions throughout the genome and therefore to observe a similar pattern when examining NA-European admixture using both genome wide loci and only *F*_ST_ outlier loci. Given the differing patterns of inferred admixture we reject this null hypothesis; the results suggest an association between European admixture and the regions of the genome most strongly contributing to differentiation of the NFL and LAB groups. This was further supported by admixture analysis of just the NFL and LAB populations based on the 1634 *F*_ST_ outliers, which showed the expected pattern of population differentiation and admixture observed previously (Figure 6C). The reanalysis of admixture with *k*=3 showed that the LAB group could be isolated by the admixture algorithm in the presence of the full European ancestry individuals, which provided evidence that the observed signal did not result from algorithm error.t A re-running of PCA using all North American samples and only the 75% of markers (n=49173) least associated with North American-European structuring resulted in a change in the pattern of genetic structure across the three North American populations (Figure S6). In a reduced PCA using only markers with the lowest p values for association with North American-European structuring, the LAB population no longer separated from the other two populations but localized between the two other populations in PC space, further suggesting that loci associated with historical European introgression have contributed to the unique genetic structure of the LAB population (Figure S7).

**Figure 6.**
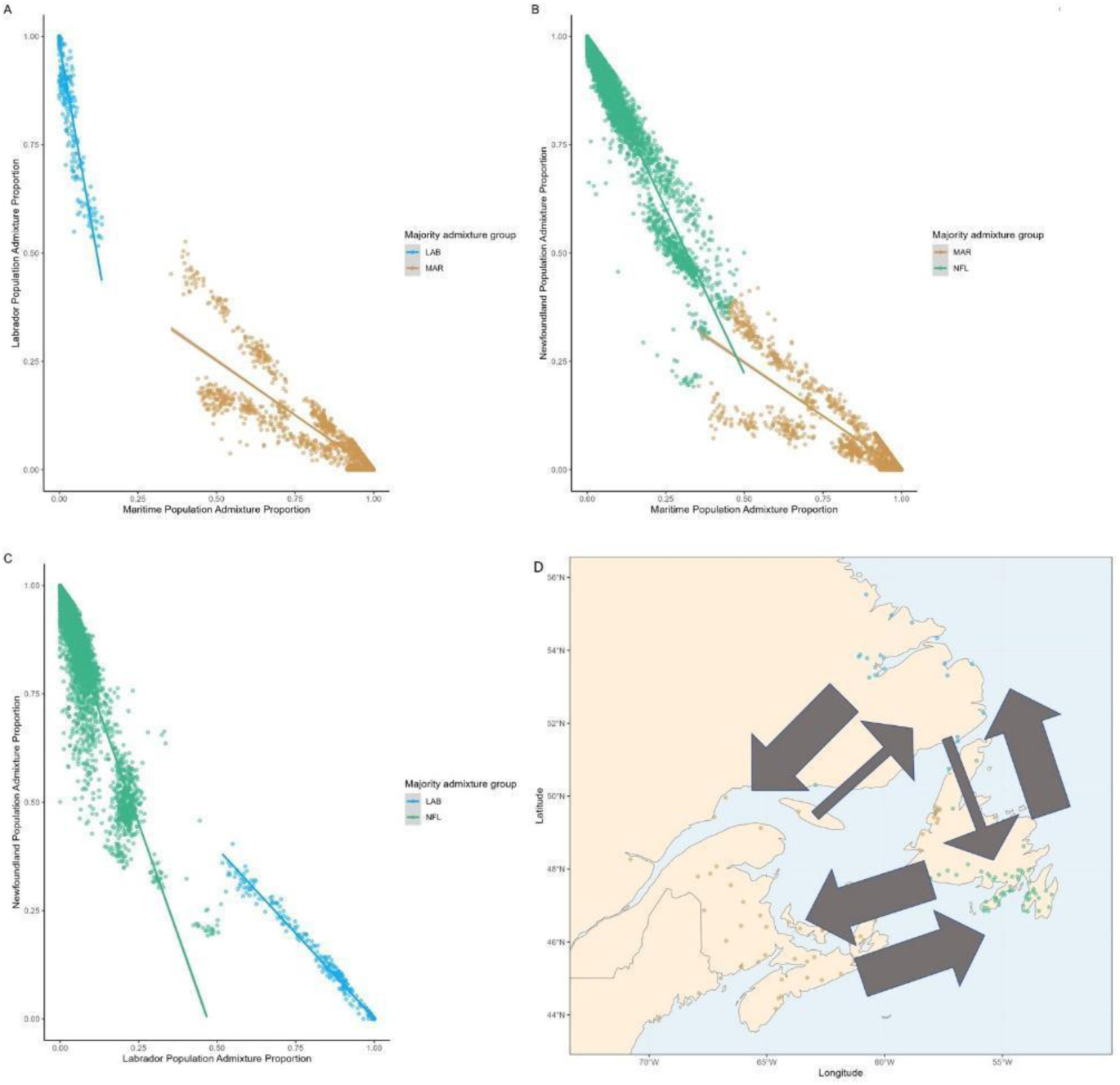
Pairwise scatter plots showing the per-individual admixture proportions of each pair of groups. The plots are limited to individuals whose majority admixture proportion is one of the two groups in the given comparison (*e.g.* for panel A comparing MAR and LAB, majority NFL individuals are omitted). Lines of best fit for each majority admixture group are shown separately. For points assigned to a given population (as indicated by colour), a diagonal scatter of points indicates the admixture from the pairwise counterpart population into the given population. A lack of diagonal points indicates that the given admixture combination is not readily occurring (e.g. for panel A: a set of MAR points are scattering along the diagonal towards the LAB, indicating admixture of the LAB population into the MAR population. No diagonal scatter of LAB points towards the MAR population is seen, suggesting a lack of admixture from MAR into LAB. Furthermore, both sets of points present a scatter of points towards position [0, 0], indicating the populations receive admixture from the NFL population, which is shown directly in the other panels). The population combinations in the plots are: A) MAR and LAB, B) MAR and NFL, C) LAB and NFL. D) A diagrammatic summary of the patterns of directional admixture suggested by the scatter plots in the proceeding three panels. Thick arrows represent evidence of directions of genetic exchange with high admixture (as indicated by diagonal scatter of points in previous scatter plots, which indicate individuals with intermediate admixture between the two groups in the comparison) and thin arrows represent directions with evidence of low admixture.

### Demographic reconstruction

The estimated maximum likelihood phylogenetic tree of populations produced using *treemix* revealed a demographic branching pattern that captured a similar population structure to the PCA and admixture analysis. The sampling locations from the three identified populations separated into distinct clades (Figure 3; Supplementary Figure S4), with the high admixture rivers (*e.g.* TRI, COD, SJQ, FLB, NAT, COR, see Table S1) localizing to the phylogenetic boundaries between the populations and displaying the highest residual fit values (Figure 3). The maximum likelihood tree suggests a basal, long-term separation of the three groups and further that the LAB population is most closely related to the NFL population, with these two groups displaying a more recent split from one another.

Examination of the geographic differences in heterozygosity (Figure 3; Table S3; Supplementary Figure S3) revealed regional differences in genomic diversity within each of the three identified groups. The heterozygosity values did not exhibit clear signatures of colonization trajectory (*e.g.* latitudinal or longitudinal gradients) and occupied a relatively narrow range across the different rivers (observed heterozygosity minimum = 0.20; maximum = 0.298; mean = 0.24; standard deviation = 0.02). For the LAB population, the highest genome wide mean heterozygosity was observed for the sampling locations within Lake Melville which is located near the latitudinal center of the population’s sampled range (Supplementary Figure S3).

The analysis of recent changes in effective population size (N_e_) revealed differences in the magnitude of N_e_ changes for the three Atlantic salmon populations (Figure 3). All three populations show evidence of dramatic declines in their recent history (past 10 generations) as well as elevated population sizes in intermediate levels of past generations (past 10-75 generations) and relatively stable N_e_ estimates, in the range of 2000-5000 individuals for their more distant past. Of the three lineages, the MAR population displayed the smallest elevation of N_e_ estimates in their intermediate past, while both the LAB and NFL populations have N_e_ estimates for this period in excess of 15000 (Figure 3).

### Gene Ontology

The 31 regions of high LD included 422 unique candidate genes for further inspection. The set of significant *F*_ST_ markers (loci with significant *F*_ST_ scores from one or more of the pairwise comparisons) overlapped with 4506 unique genes. The significant loci from the PCA overlapped with a series of 278 genes. There were no instances of a gene being identified in all three analyses (*e.g.* containing significant loci from the PCA and *F*_ST_ analyses and also falling within a window of high LD). However, 57 unique genes were associated with the significance peaks (SNPs or LD regions) from two of the three analyses (Table S4). These genes spanned 24 of the 29 Atlantic salmon chromosomes (chromosomes 12, 16, 20, 24, and 29 were not represented). The largest concentration of identified genes were seen on chromosomes 1 and 4, where 8 genes of interest were each identified. Three specific genes of interest identified in previous studies: *vgll3*, *six6*, and *MHCII* were not seen within any of the sets of candidate genes here identified. The Entrez GeneIDs for the 57 genes of potential interest identified in the cross referencing of analyses were then examined using ShinyGo. Seven pathways were identified as significantly enriched (FDR P < 0.05) (Table S5).

## DISCUSSION

Pleistocene glaciations and subsequent post-glacial recolonization in northern environments have been significant determinants of diversity impacting both adaptation to the environment and future responses of populations and species to climate change (e.g., Rendon-Anaya *et al*. 2021; Luqman *et al*. 2023). Here we provide the most extensive genomic and geographic evaluation of diversity of Atlantic salmon throughout the Northwest Atlantic range to date. Our results suggest three genetically and geographically distinct phylogeographic groups with high genome-wide divergence coupled with admixture near their geographic boundaries. In addition to regional secondary contact within North America, low levels of European admixture were detected within the Newfoundland and Labrador sampling locations, strongly suggesting historical secondary contact with migrants from Europe. Genome regions associated with isolation of the Labrador population also contained higher proportions of European admixture than the genome as a whole, suggesting a role of European introgression in population divergence. Overall, these results suggest that contemporary Atlantic salmon population structure has been influenced by neutral evolutionary processes (*e.g.* drift in allopatry) and admixture of lineages in regions of secondary contact on two different scales: intra-and inter-continental. This work provides the most comprehensive examination of the Atlantic salmon genomic landscape within northeastern North America to date, building upon previous trans-Atlantic (Moore *et al*. 2014; Rougemont & Bernatchez 2018; Lehnert *et al*. 2020) and regional studies (Sylvester *et al*. 2018; Wellband *et al*. 2019). The detailed description of the genomic landscape also provides novel insights into broadscale population structure that can be considered in future management and conservation efforts in this species.

### Demographic patterns and post-glacial secondary contact

Our evidence of large-scale genetic structure in Atlantic salmon is suggestive of recolonization of the region by at least two distinct lineages likely recolonizing following the most recent deglaciation of northeastern North America, which occurred ∼14-11 Kya (Shaw *et al*. 2006). We propose that a western glacial lineage is represented by the Maritime group and the second eastern lineage by the Newfoundland and Labrador groups. While this represents the first strong genomic evidence of two glacial lineages in North America, this aligns with previous descriptions of inferred refugia for other species in the region, which posit the existence of extensive southern glacial refugia (Taberlet *et al*. 1998; Bernatchez & Wilson 1998; Hewitt 2004; Maggs *et al*. 2008). The areas of high contemporary genetic admixture between the western and eastern lineages align with the Gulf of St. Lawrence Ice Flow (Shaw *et al*. 2006), further suggesting a role glaciation in shaping the colonization patterns of the lineages. The two lineages are also concordant with previous examination of North American Atlantic salmon population structure by Rougemont & Bernatchez (2018) that identified the existence of two major North American groups exhibiting north-south clustering. However, our identification of three extant groups in North America builds upon previous descriptions and suggests the contemporary population structure cannot be attributed solely to patterns of post glacial recolonization from two refugia. At first appearances, interpretation of the Labrador population as stemming from a third, periglacial refugium is enticing (and remains an alternative hypothesis that we do not have sufficient evidence to reject). For other species, small periglacial refugia at more northern latitudes have been described and are proposed as regions where small, isolated populations could potentially have persisted. (Petit *et al*. 2003; Provan & Bennet 2008). However, based on the genomic structure, phylogenetic analysis, and the patterns of genetic differentiation we present, it appears more likely that the Labrador and Newfoundland groups are derived from a common ancestral lineage (*e.g.* a lineage synonymous to the more eastern of the two major groups described by Rougemont & Bernatchez (2018)) that colonized the entirety of the Eastern coasts of Newfoundland and Labrador from the south following the Pleistocene deglaciation. This scenario of population origin and structure has interesting parallels with the patterns of post-glacial recolonization of the Baltic by Atlantic salmon, which is proposed to have involved two lineages derived from separate refugia (Verspoor *et al*. 1999; Consuegra *et al*. 2002; King *et al*. 2007; Finnegan *et al*. 2013; Rougemont & Bernatchez 2018), one of which has subdivided into two groups following northward colonization by a southern lineage (Biere *et al*. 2011; Rougemont & Bernatchez 2018).

In addition to secondary contact between northwest Atlantic refugia, our results support previous work indicating secondary contact with migrants from Europe, particularly in the easternmost regions, that is estimated to have begun ∼13,400 years ago (95% CI = 1,800–41,380; Rougemont & Bernatchez 2018). The signal of European admixture along the eastern range of sampling locations shows that low levels (2- 4%) of European admixture persist in individuals from these regions and may contribute to observed differences with the more western MAR group, where little to no evidence of European admixture is seen. The overrepresentation of regions of the genome displaying European admixture in the differentiation of the two eastern groups (Newfoundland and Labrador) may be the result of the interaction of the two signals of secondary contact (*e.g.* admixture both between the North American glacial lineages and between the European and eastern North American lineage). Following deglaciation, migration and admixture between the NFL and MAR groups may have reduced the magnitude of European admixture within parts of the NFL group and thereby contributing to its differentiation from the more northern LAB group where lower levels of admixture from the west are observed and the European admixture signal may therefore be less diluted as a result. The genomic composition of the NFL group may therefore have been shaped by the interaction of two secondary contact events, one with the MAR glacial lineage from the west and a second with the European lineage from the east. This is in line with previous conclusions of Rougemont and Bernatchez (2018) who concluded that the demographic history of Atlantic salmon in North America was shaped by multiple secondary contacts both between and within the North American and European continents. Admittedly, evidence is not sufficient to implicate European introgression as a main driver of population structure (Watson *et al*. 2022), but there is a detectable signal associated with extant population structure and signals of differentiation between the NFL and LAB groups. One putative explanation for the European-associated differences between the NFL and LAB groups could be secondary contact hailing from different European source populations or different colonization events from the same source. Although the exact cause cannot be stated with certainty, these regions of the genome are likely to some extent a factor contributing to the population structure, along with other influences such as climate associated adaptation that will require further characterization in future studies. Additionally, the similarity in estimated timing of European secondary contact and proposed recolonization from a southern refugium following deglaciation (∼14-11 Kya Shaw *et al*. 2006) suggest that European secondary contact co-occurred with colonization of the region (*e.g.* as habitat became available, it may have been simultaneously colonized by more proximal individuals originating from southern refugia and in lesser numbers by individuals from more distant European sources).

Another possible cause of the differentiation of the LAB and NFL groups is environment-associated divergence. Within Europe, environment-associated adaptive divergence has been proposed as a mechanism shaping extant population structure following post-glacial recolonization, with local adaptation to a marginal environment proposed as the causative agent of the genetic differentiation observed between populations of fish in the Baltic and North Seas (Bierne *et al*. 2011). The LAB population encompasses the northern limit of our sampling range and extends to near the known northern edge of the range of Atlantic salmon (DFO 2018; Sylvester *et al*. 2018). Therefore, environment-associated selective pressures that are unique to, or magnified at, the northern extremes of the range may be contributing to divergence of these lineages. The examination of pairwise genomic differentiation across the populations provided minimal distinct peaks of differentiation, which suggests neutral, as opposed to adaptive, genetic differences between the populations with the most prominent peaks observed consistently associated with the LAB group. Similarly, the search for genes of interest within regions identified in the differentiation analysis contained genes associated with several specific pathways and failed to provide a strong signal of a specific adaptive mechanism. The gene search also did not reveal any association with *vgll3 (*Ayllon *et al*. 2015), *six6* (Moustakas-Verho *et al*. 2020), or *MHCII* (Kjøglum *et al*. 2008), all of which have previously identified as associated with adaptive evolution in Atlantic salmon. It is possible that the hierarchical nature of salmon populations and previously characterized regional genomic structuring resulting from the species’ strong homing behaviour (Dodson *et al*. 1998; Fraser *et al*. 2011; Moore *et al*. 2014; Wellband *et al*. 2018) has caused local adaptive differences to be overlooked due to a genomic background of elevated population structure (Waples *et al*. 2022). Future work is thus required to demonstrate the presence of adaptive divergence to regional environmental variation, and the genes and associated pathways involved at this spatial scale.

### Comparison to demographic histories of other marine species in the northwest Atlantic

The inferred glacial lineages of Atlantic salmon in Atlantic Canada add to the number of species with described recolonization patterns in the Atlantic Canada region (McCusker & Bentzen 2010; Brunner *et al*. 2001; Moore *et al*. 2015; Bringlow *et al. in review*). Most notably, there are similarities to the characterized genetic structure of rainbow smelt (*Osmerus mordax*) from coastal Newfoundland, for which genetic data has supported the hypothesis that contemporary spatial structure reflects historical landscape isolation resulting from glacial cycles and maintained by low dispersal and selective processes; broader patterns observed in smelt though may suggest increased complexity (Bernatchez 1997; Bradbury *et al*. 2011. Genetic analysis of rainbow smelt also suggest two glacial lineages present throughout Atlantic Canada (Bernatchez 1997). The multi-refuge recolonization described for Atlantic salmon here is also highly similar to the described recolonization of the region by Arctic charr, a closely related species (Brunner *et al*. 2001; Moore *et al*. 2015). For Arctic charr, two genetically distinct lineages have been hypothesized to have been present in Atlantic Canada, stemming an Atlantic, and an Acadian refugium, for which the distribution of lineages broadly coincide with the western MAR and eastern NFL and LAB salmon lineages. The similarity of the Arctic charr and Atlantic salmon colonization patterns is especially interesting given the variable recolonization patterns observed across other species in the region, such as species of wolffish, which appear to have undergone post glacial expansion from a single location (McCusker & Bentzen 2010). It is possible the similar recolonization patterns of Arctic charr and Atlantic salmon are due to the existence of southern refugia that presented habitats suitable for both salmonid species to persist in during the last ice age and that they expanded their ranges in similar fashions as glaciers receded. Although, given differences in aspects of life history (*e.g.* freshwater Arctic charr populations (Kess *et al*. 2021)), phylogeography (*e.g.* more northern range of Arctic charr (Power *et al*. 2005)) and demographic history (*e.g.* the role of European migrants in shaping Atlantic salmon population structure (Lehnert *et al*. 2019)), these similarities may be coincidental, or the result of similar evolutionary responses to environmental variation across the common geographic range (Stanley *et al*. 2018).

### Additional admixture parallels

There is also evidence of a similar post-glacial recolonization history for Atlantic salmon within their European range. As previously mentioned, inferred post-glacial recolonization of the Baltic Sea by Atlantic salmon, is proposed to have involved two lineages derived from separate refugia (Verspoor *et al*. 1999; Koljonen *et al*. 1999; Consuegra *et al*. 2002; King *et al*. 2007; Finnegan *et al*. 2013; Rougemont & Bernatchez 2018), one of which has subdivided into two groups following northward colonization by a southern lineage and resulting in three contemporary geographic groups (Asplund *et al*. 2004; Biere *et al*. 2011; Rougemont & Bernatchez 2018). Some of the finer scale patterns here described have European parallels as well, such as evidence of localized North American introgression into the Barents Sea populations of northern Europe (Makhrov *et al*. 2005); salmon in this region are also considered a different phylogeographic group but may be placed within the same glacial lineages as other more southern populations (Wennevik *et al*. 2019). In addition, as observed here for North American populations, there are documented transition zones with admixture between the phylogeographic groups of Europe (Wennevik *et al*. 2019).

It has not escaped our notice that the patterns of European secondary contact into North American Atlantic salmon bear a strong resemblance to the patterns seen in other species, including the introgression of Neanderthal (*Homo neanderthalensis*) DNA into human (*Homo sapiens*) populations. Approximately 2–4% of genetic material in human populations outside Africa is derived from Neanderthals who interbred with anatomically modern humans (Harris & Nielsen 2016; Dannemann & Racimo 2018). Evidence suggests that multiple episodes of interbreeding between Neanderthal and modern humans and of multiple episodes of gene flow into both European and East Asian populations have occurred (Vilanea & Schraiber 2019), which is similar to Atlantic salmon European secondary contact patterns discussed by Lehnert *et al*. (2019) and Rougemont & Bernatchez (2018) and further supported by our findings.

## Conclusion

Our analyses of the genomic structure of Atlantic salmon populations across most of the species’ range in North America revealed three genetically distinct population groups that occupy distinct portions of the contemporary range. Exploration of the origins of the three population groups suggests post-glacial recolonization of the region by glacial lineages originating from two refugia. Salmon from a western refugium recolonized southern Atlantic Canada and northern New England, whereas those from an eastern refugium recolonized insular Newfoundland and mainland Labrador. We further conclude that during or following postglacial dispersal the eastern lineage experienced secondary contact with salmon of European origin, and that variation in the resulting admixture contributed to the development of two genetically distinct population groups in Newfoundland and Labrador, respectively, as well as their differentiation from the population group derived from the western lineage. Areas of genetic admixture in border areas of the western and eastern lineages align with the Gulf of St. Lawrence Ice Flow, suggesting that the glacier shaped early dispersal of the lineages, and its subsequent recession enabled localized secondary contact between the lineages. Therefore, spatially variable introgression between the North American glacial lineages and between the eastern lineage and European migrants appears to have influenced the contemporary population genomic structure of Atlantic salmon in North America. Future work should examine in greater detail how historical contingency and environmentally driven adaptive variation have shaped genomic variation and demographic patterns in Atlantic salmon, and how genomes and populations are likely to respond in the face of rapid climate change.

## DATA AVAILABILITY STATEMENT

Data used in these analyses are publicly available and were generated in previous studies (Bradbury et al., 2018, 2022; Lehnert et al., 2019; Nugent *et al*. 2023).

## SUPPORTING INFORMATION

Supplementary_tables_and_figures.pdf – A file containing the supplementary tables and figures for the manuscript.

## ACKNOWLEDGEMENTS

The authors would like to thank staff of Fisheries and Oceans Newfoundland and Labrador Salmonids Section for assistance with sample collection. Funding was provided through the Program for Aquaculture Regulatory Research of Fisheries and Oceans Canada, the Genomics Research and Development Initiative of Canada, and the National Sciences and Engineering Research Council of Canada.

