## Supplementary_tables_and_figures for "Post-glacial recolonization and multiple scales of secondary contact contribute to contemporary Atlantic salmon (*Salmo salar*) genomic variation in North America"

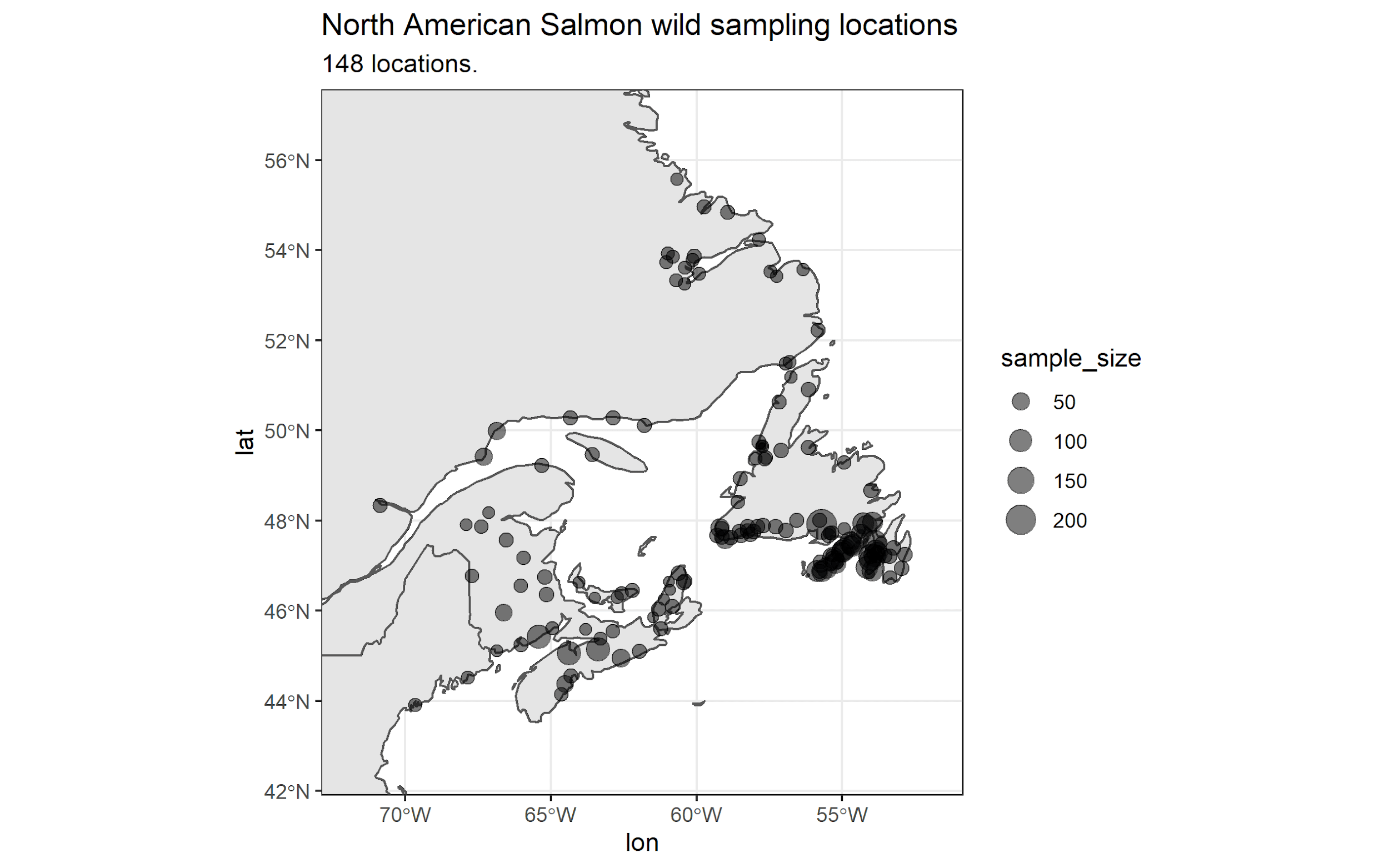

Figure S1. Map of the 148 Atlantic salmon sampling locations. The size of the points is reflective of the number of samples from each location that were genotyped with the SNP array.

A)

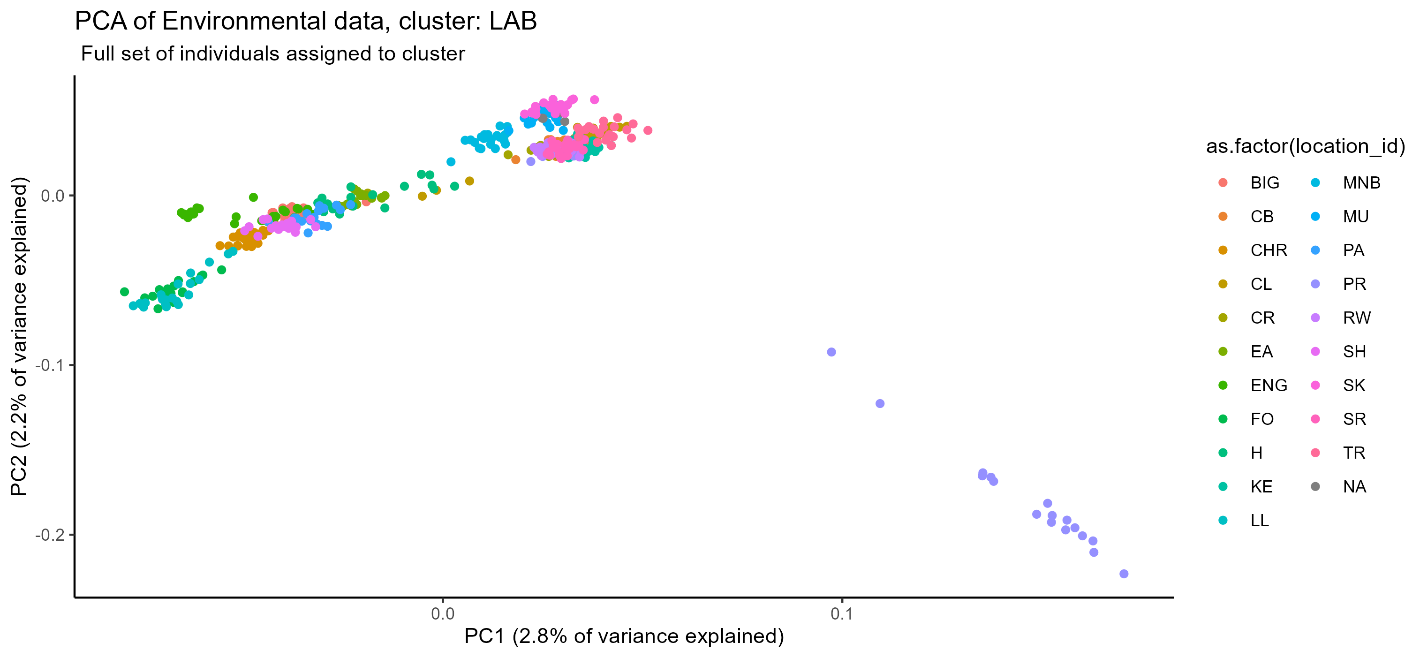

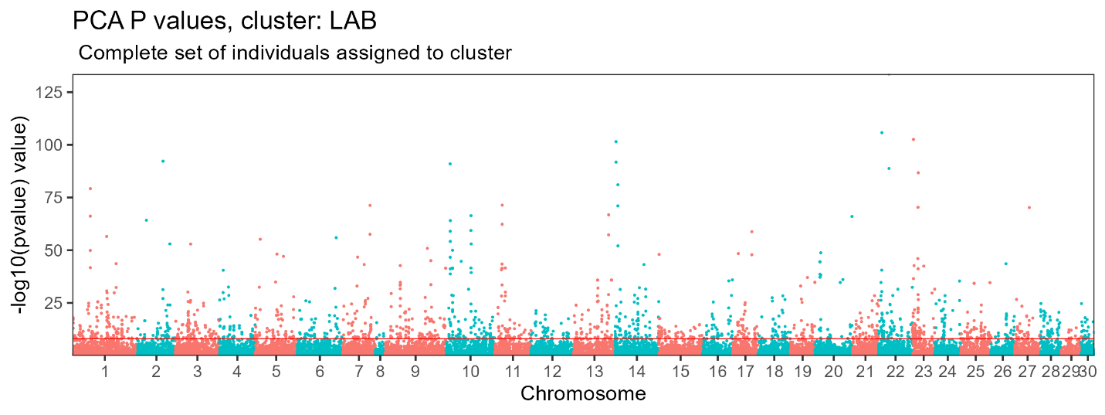

B)

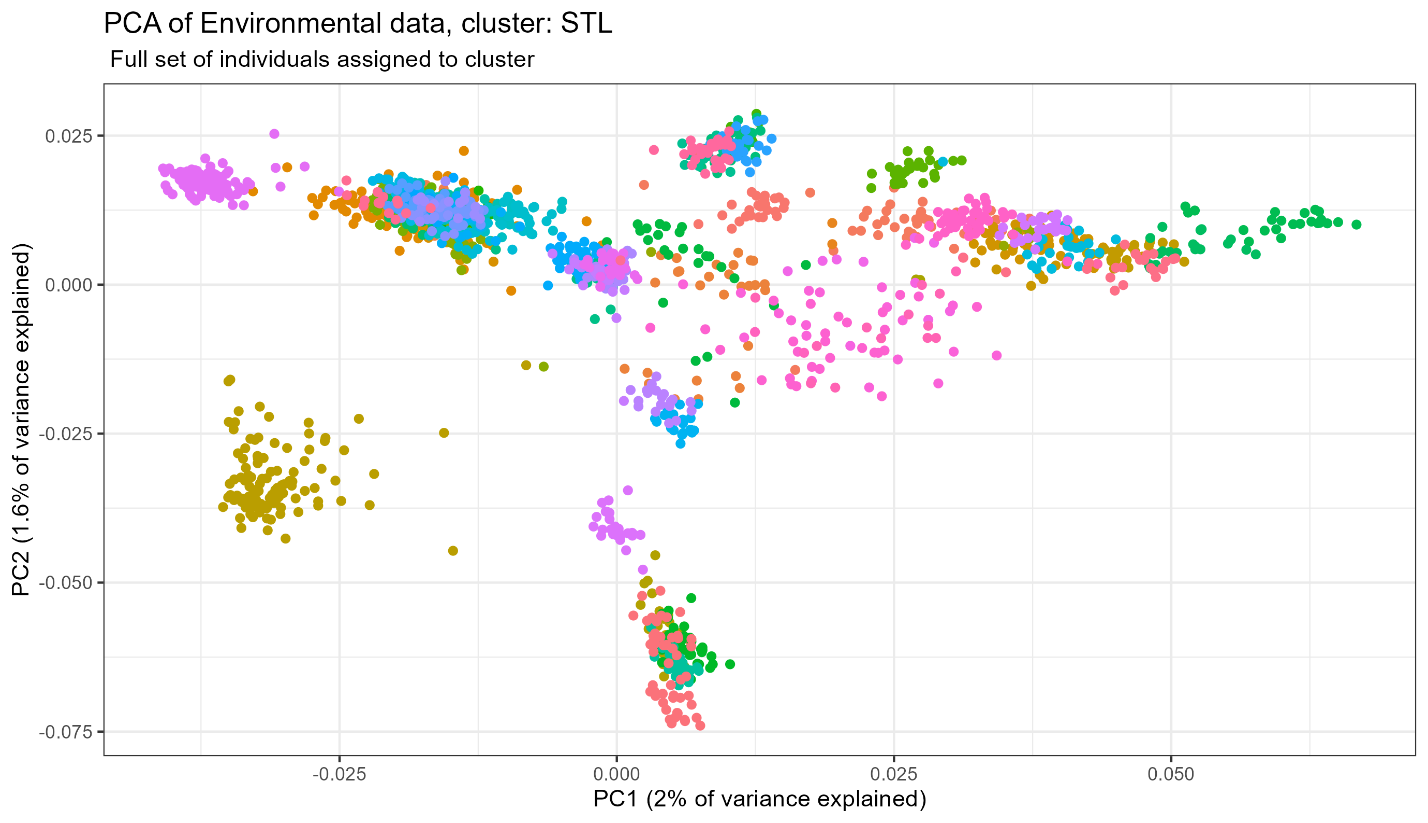

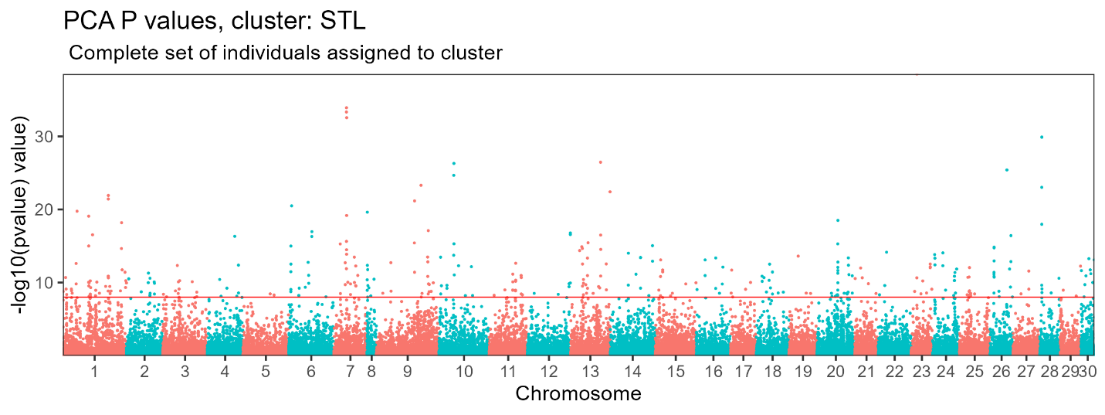

C)

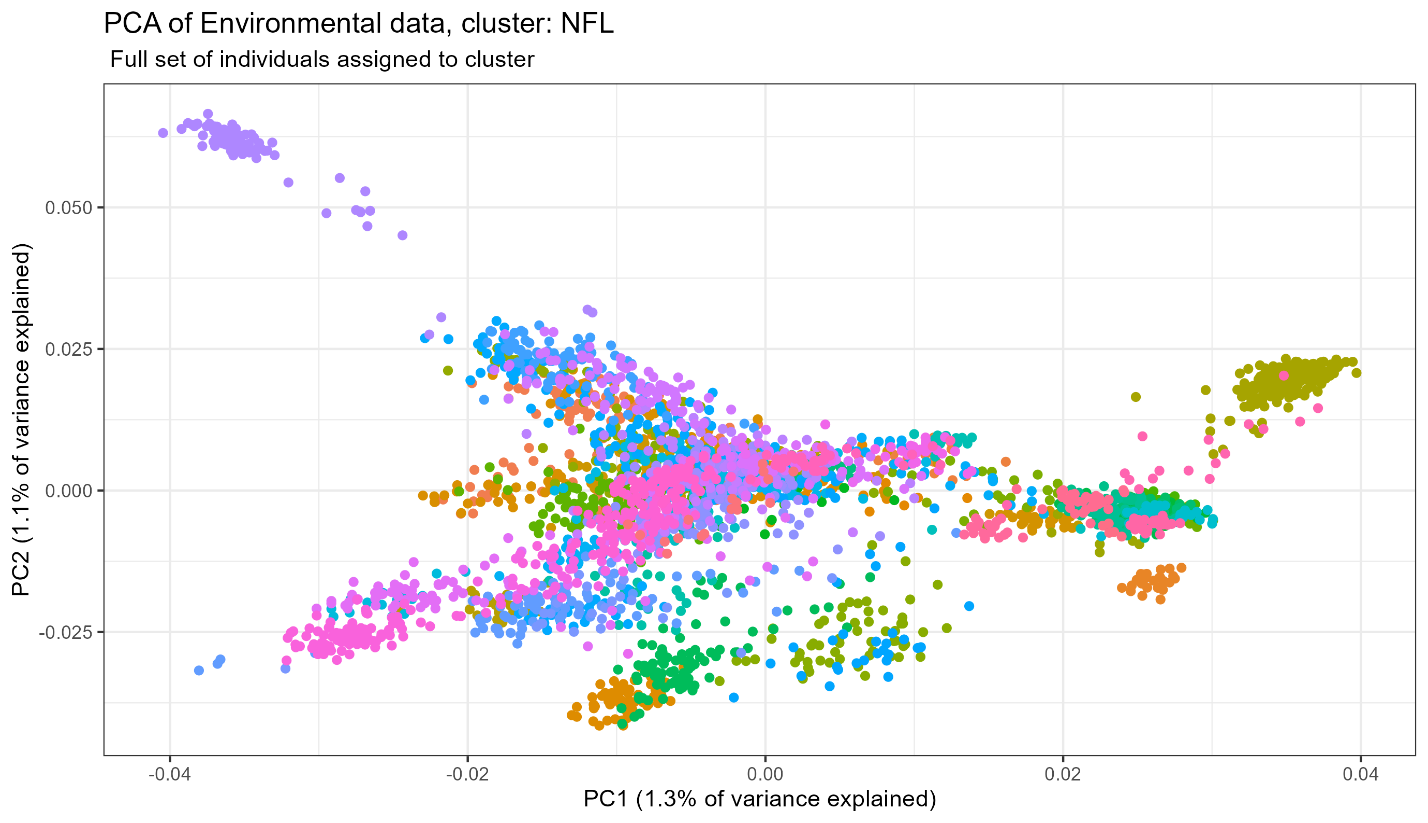

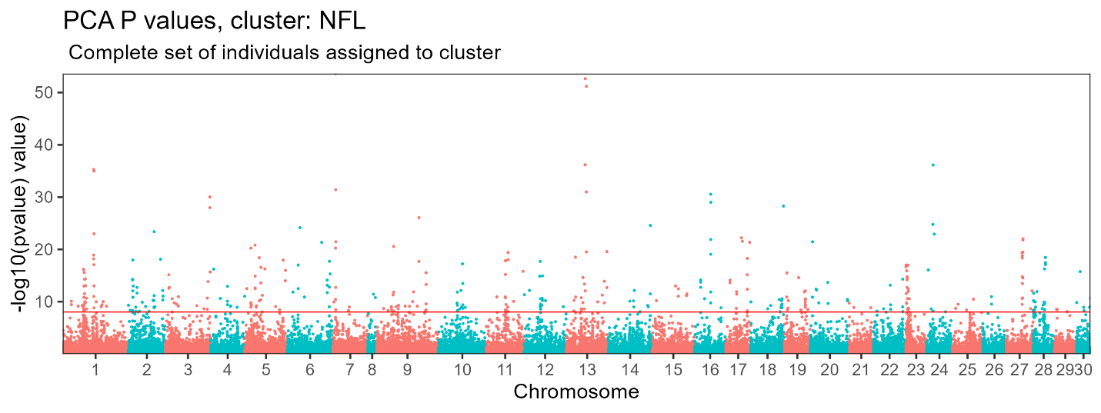

Figure S2. Scatter plots of Principal Components (PCs) of genetic variation and Manhattan plots showing the P values of each marker with the PC values, for each of the three identified populations: A) Labrador, B) Maritimes, and C) Newfoundland. Points are coloured by location of origin.

A)

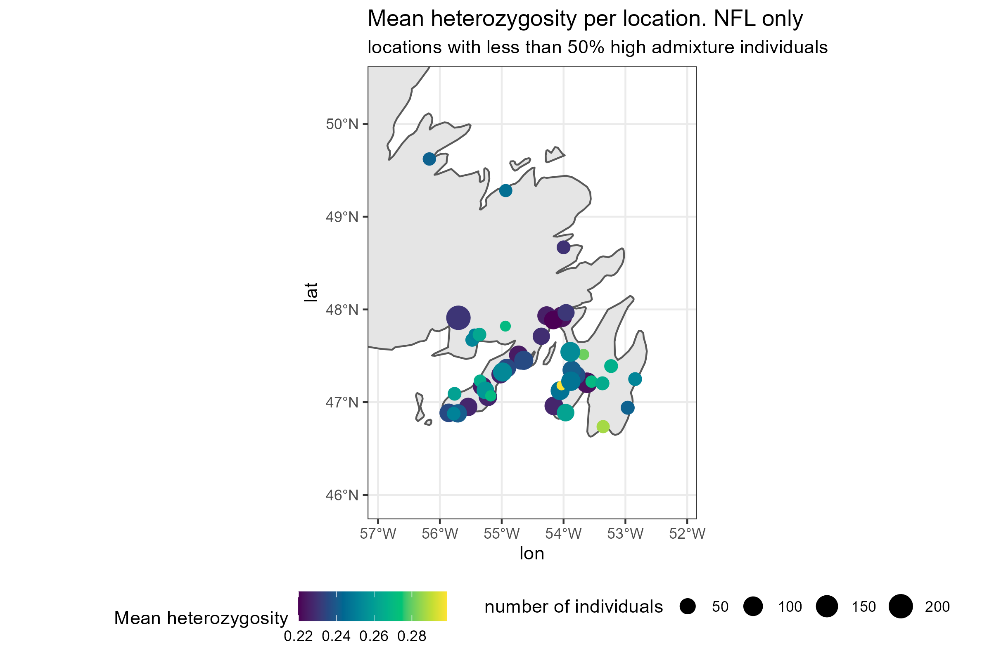

B)

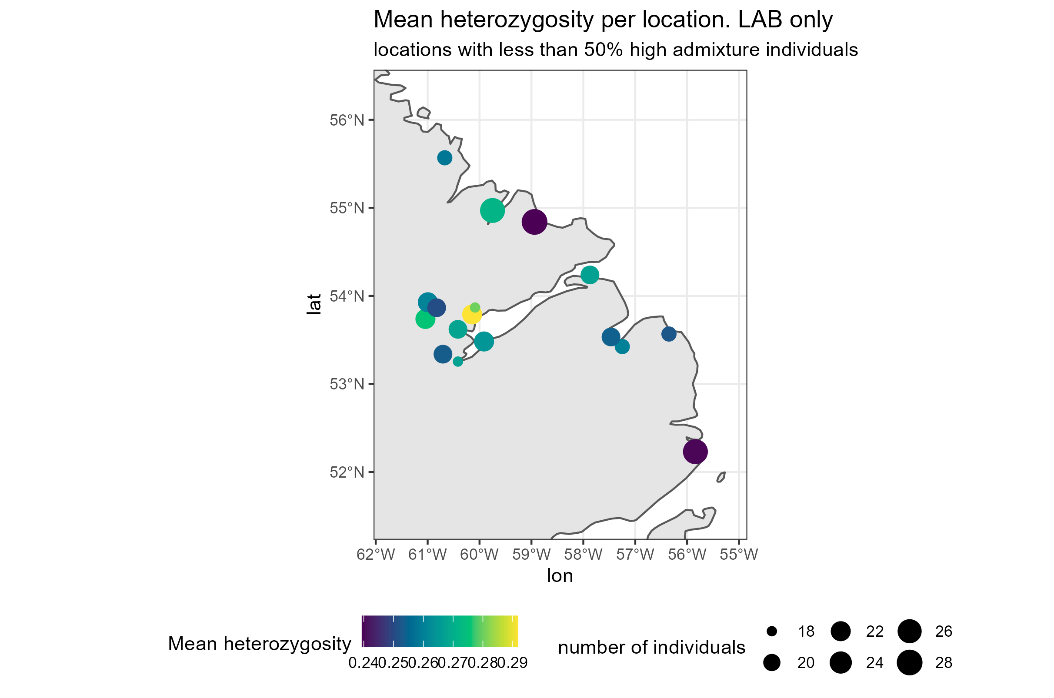

C)

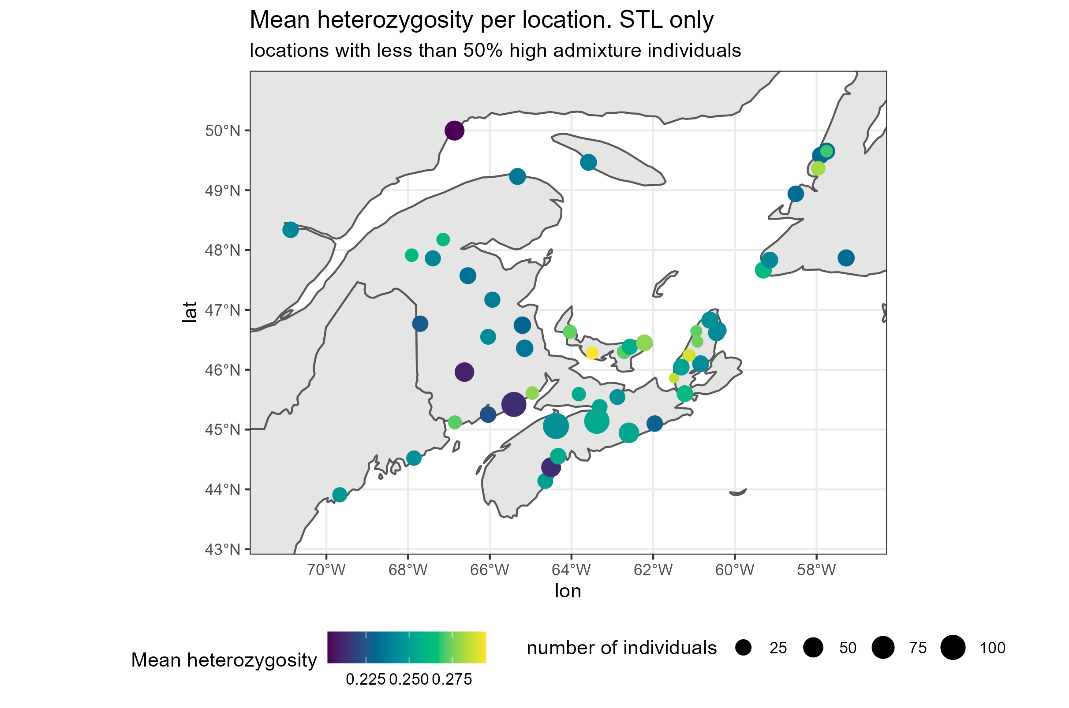

Figure S3. Per-location mean heterozygosity for all sampling locations. The colour of the points indicate the genome wide mean heterozygosity for the sampling location and the size of the points indicate the sample size for the given location . The map is broken down by population and only location with <50% high admixture individuals are shown. Populations shown are: A) Newfoundland (NFL) B) Labrador (LAB) C) Maritimes (MAR)

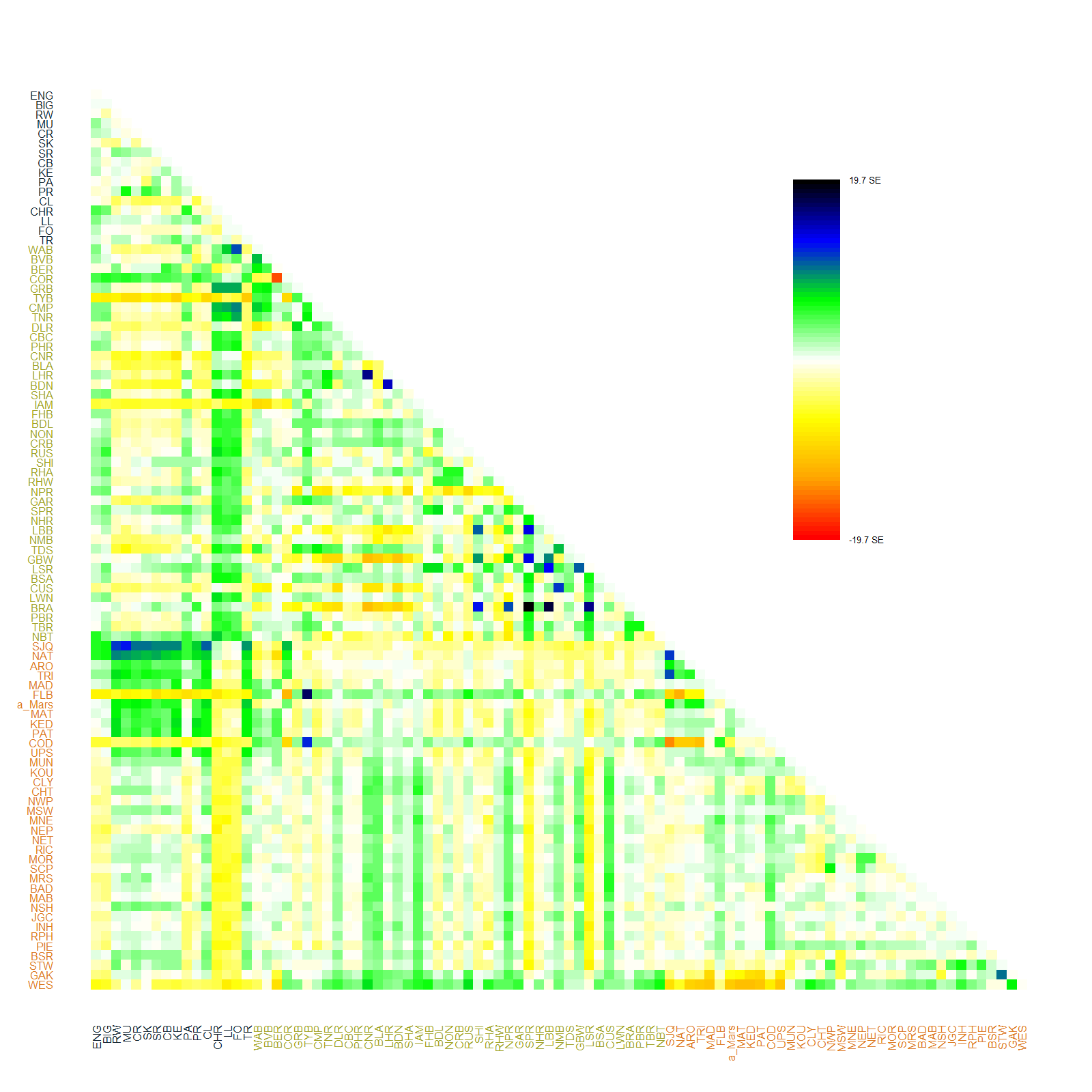

Figure S4. Matrix of the residual values of all the pairwise sampling location comparisons conducted in the *treemix* analysis.

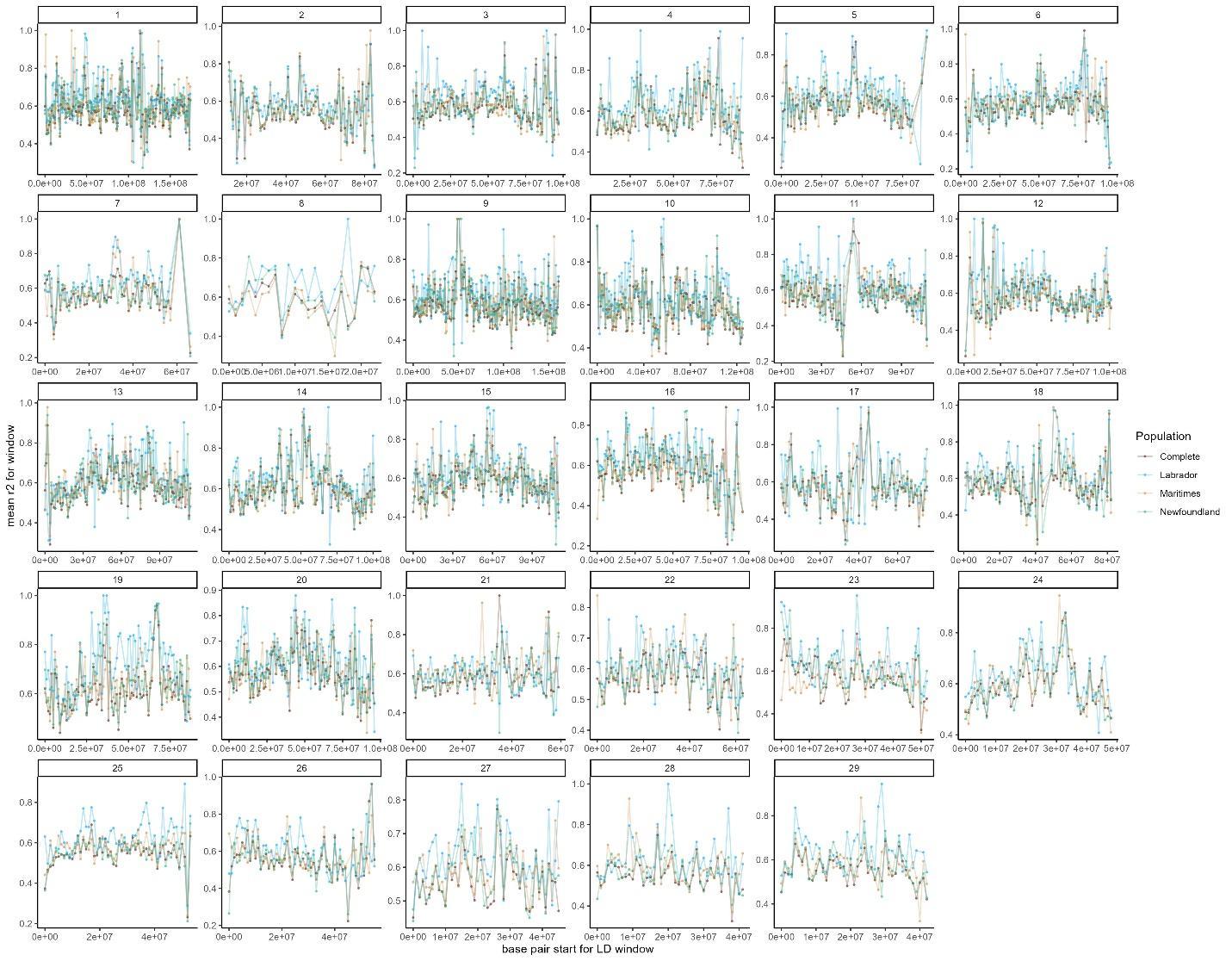

Figure S5. Per-chromosome windowed (1mb) mean Linkage Disequilibrium (LD) for each of the three identified Atlantic salmon populations and the complete set of individuals.

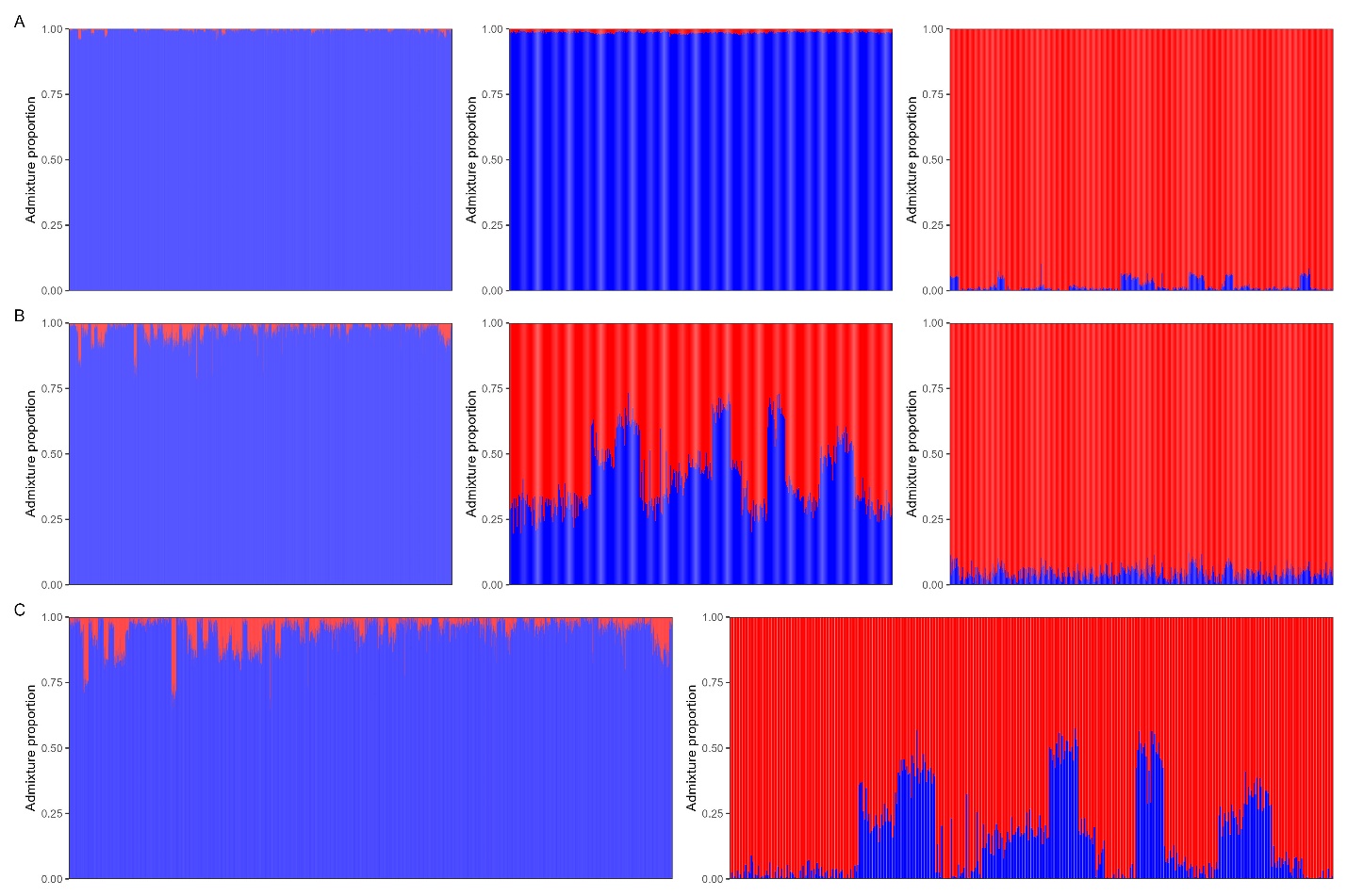

Figure S6. A) Admixture proportion plot (with *k* = 2) based on genome wide SNP set for three sets of individuals: NFL (left), LAB (middle), and Norwegian wild (right). B) Admixture proportion plot (with *k* = 2) based on the Fst (n= 1634 SNPs) outlier SNPs from the LAB-NFL comparison. Admixture proportions shown for three sets of individuals: NFL (left), LAB (middle), and Norwegian wild (right) C) Admixture proportion plot (with *k* = 2) based on the *F*_ST_ outlier SNPs (n= 1634 SNPs) from the LAB-NFL comparison and two sets of individuals: NFL (left), LAB (right).

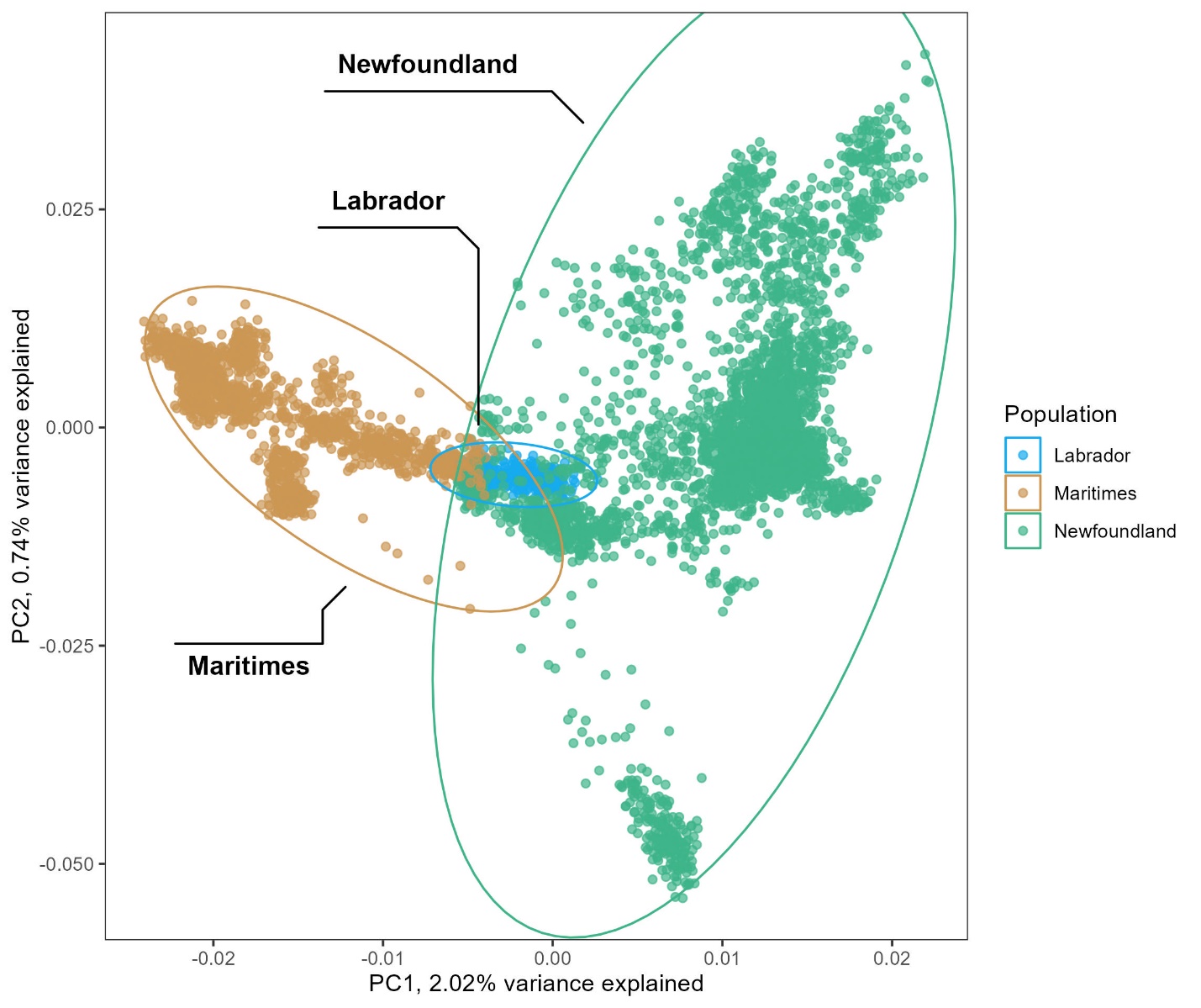

Figure S7. Scatter plot of Principal Components (PCs) of reduced summarization of genetic variation for 5455 Atlantic salmon individuals sampled. This PCA was conducted using only the 49173 (75% of polymorphic markers) displaying the lowest p values for association with North American-European structuring (*e.g.* Figure 2A of Bradbury *et al.* 2022). The coloured ellipses denote the genetic populations identified through *k*-means clustering of whole genome PCs and the cluster to which individuals were assigned. Through comparison to Figure 1B. we can infer that without the 25% of loci most significantly associated with North American-European genetic structure, the Labrador population is no longer strongly separated from the other two populations.

TABLE S1. Per location summary statistics.

|  | | | | | K-means clustering | | | | Admixture | | Heterozygosity | |
| --- | --- | --- | --- | --- | --- | --- | --- | --- | --- | --- | --- | --- |
| ID | Location name | Latitude | Longitude | Sample number | Majority assigned cluster | LAB count | NFL count | MAR count | Are majority of individuals admixed? | Frequency of admixed individuals | Mean observed | Mean expected |
| TR | Trout River, Labrador | 49.65 | -57.76 | 22 | LAB | 22 | 0 | 0 | FALSE | 0.00 | 0.271 | 0.267 |
| SR | Susan River, Labrador | 53.74 | -61.04 | 22 | LAB | 22 | 0 | 0 | FALSE | 0.00 | 0.276 | 0.270 |
| CHR | St Charles River, Labrador | 52.23 | -55.84 | 27 | LAB | 27 | 0 | 0 | FALSE | 0.04 | 0.240 | 0.235 |
| SK | Sebaskachu River, Labrador | 53.79 | -60.14 | 22 | LAB | 22 | 0 | 0 | FALSE | 0.00 | 0.292 | 0.285 |
| SH | Sand Hill River, Labrador | 53.57 | -56.35 | 19 | LAB | 19 | 0 | 0 | FALSE | 0.00 | 0.252 | 0.248 |
| RW | Red Wine River, Labrador | 53.93 | -61.00 | 22 | LAB | 22 | 0 | 0 | FALSE | 0.00 | 0.261 | 0.258 |
| PR | Peters River, Labrador | 53.34 | -60.71 | 21 | LAB | 21 | 0 | 0 | FALSE | 0.00 | 0.253 | 0.256 |
| PA | Paradise River, Labrador | 53.42 | -57.25 | 19 | LAB | 19 | 0 | 0 | FALSE | 0.00 | 0.260 | 0.257 |
| MU | Mulligan River, Labrador | 53.87 | -60.09 | 18 | LAB | 18 | 0 | 0 | FALSE | 0.00 | 0.280 | 0.274 |
| MNB | Main Brook, Labrador | 54.24 | -57.87 | 21 | LAB | 21 | 0 | 0 | FALSE | 0.00 | 0.267 | 0.262 |
| LL | L'anseau Loup River, Labrador | 51.53 | -56.82 | 22 | LAB | 20 | 2 | 0 | TRUE | 0.91 | 0.251 | 0.247 |
| KE | Kenamu River, Labrador | 53.48 | -59.91 | 22 | LAB | 22 | 0 | 0 | FALSE | 0.00 | 0.264 | 0.260 |
| H | Hunt River, Labrador | 55.57 | -60.67 | 19 | LAB | 19 | 0 | 0 | FALSE | 0.00 | 0.258 | 0.262 |
| FO | Forteau River, Labrador | 51.48 | -56.94 | 21 | LAB | 21 | 0 | 0 | TRUE | 1.00 | 0.246 | 0.246 |
| ENG | English River, Labrador | 54.97 | -59.75 | 27 | LAB | 27 | 0 | 0 | FALSE | 0.00 | 0.272 | 0.264 |
| EA | Eagle River, Labrador | 53.53 | -57.47 | 21 | LAB | 21 | 0 | 0 | FALSE | 0.00 | 0.254 | 0.253 |
| CR | Crooked River, Labrador | 53.87 | -60.83 | 21 | LAB | 21 | 0 | 0 | FALSE | 0.00 | 0.251 | 0.255 |
| CL | Caroline River, Labrador | 53.25 | -60.42 | 18 | LAB | 18 | 0 | 0 | FALSE | 0.00 | 0.267 | 0.272 |
| CB | Cape Caribou, Labrador | 53.62 | -60.42 | 21 | LAB | 21 | 0 | 0 | FALSE | 0.00 | 0.268 | 0.264 |
| BIG | Big River, Labrador | 54.84 | -58.94 | 28 | LAB | 28 | 0 | 0 | FALSE | 0.00 | 0.239 | 0.244 |
| COR | Corneille, Quebec | 50.28 | -62.88 | 28 | NFL | 0 | 23 | 5 | TRUE | 1.00 | 0.240 | 0.245 |
| NAT | Natashquan, Quebec | 50.12 | -61.80 | 28 | MAR | 0 | 2 | 26 | TRUE | 0.93 | 0.212 | 0.220 |
| TRI | Rivire de la Trinite, Quebec | 49.42 | -67.30 | 49 | MAR | 0 | 0 | 49 | TRUE | 0.88 | 0.217 | 0.217 |
| SJQ | Saint-Jean, Quebec | 50.28 | -64.33 | 28 | MAR | 0 | 0 | 28 | TRUE | 1.00 | 0.230 | 0.228 |
| NHR | North Harbour River, Newfoundland | 47.21 | -53.62 | 117 | NFL | 0 | 117 | 0 | FALSE | 0.02 | 0.223 | 0.230 |
| BER | Big East, Newfoundland | 50.63 | -57.17 | 27 | NFL | 0 | 27 | 0 | TRUE | 1.00 | 0.256 | 0.252 |
| CRB | Cape Roger Brook | 47.44 | -54.69 | 56 | NFL | 0 | 56 | 0 | FALSE | 0.00 | 0.236 | 0.238 |
| ARO | Rivire Aux Rochers, Quebec | 50.00 | -66.86 | 48 | MAR | 0 | 1 | 47 | FALSE | 0.46 | 0.203 | 0.205 |
| SIM | Simms Brook, Newfoundland | 47.67 | -55.48 | 28 | NFL | 0 | 28 | 0 | FALSE | 0.11 | 0.253 | 0.239 |
| BVB | Beaver Brook, Newfoundland | 50.90 | -56.15 | 29 | NFL | 0 | 29 | 0 | TRUE | 0.97 | 0.237 | 0.233 |
| WBR | White Bear River, Newfoundland | 47.87 | -57.28 | 30 | NFL | 0 | 30 | 0 | TRUE | 0.97 | 0.232 | 0.227 |
| WAB | Western Arm, Newfoundland | 51.19 | -56.76 | 18 | NFL | 0 | 18 | 0 | TRUE | 1.00 | 0.261 | 0.254 |
| DLR | Dollards Brook, Newfoundland | 48.02 | -56.57 | 26 | NFL | 0 | 26 | 0 | TRUE | 0.58 | 0.235 | 0.230 |
| GRR | Grey River, Newfoundland | 47.77 | -56.93 | 30 | NFL | 0 | 29 | 1 | TRUE | 1.00 | 0.231 | 0.226 |
| IAM | Isle aux Morts River, Newfoundland | 47.59 | -59.01 | 57 | NFL | 0 | 51 | 6 | TRUE | 0.98 | 0.228 | 0.226 |
| GND | Grandy Brook, Newfoundland | 47.89 | -57.72 | 29 | NFL | 0 | 27 | 2 | TRUE | 1.00 | 0.233 | 0.228 |
| CBR | Couteau Brook, Newfoundland | 47.75 | -58.03 | 30 | NFL | 0 | 30 | 0 | TRUE | 1.00 | 0.230 | 0.225 |
| GBR | Grand Bay, Newfoundland | 47.63 | -59.13 | 30 | NFL | 0 | 30 | 0 | TRUE | 1.00 | 0.232 | 0.226 |
| GNB | Grandy Brook, Newfoundland | 47.63 | -58.83 | 30 | NFL | 0 | 30 | 0 | TRUE | 1.00 | 0.235 | 0.227 |
| EBB | East Bay Brook, Newfoundland | 47.77 | -58.25 | 29 | NFL | 0 | 29 | 0 | TRUE | 1.00 | 0.256 | 0.248 |
| BSA | Big Salmonier Brook, Newfoundland | 47.06 | -55.22 | 81 | NFL | 0 | 81 | 0 | FALSE | 0.00 | 0.227 | 0.231 |
| CCR | Cinq Cerf, Newfoundland | 47.70 | -58.15 | 30 | NFL | 0 | 30 | 0 | TRUE | 1.00 | 0.226 | 0.222 |
| LPR | La Poile, Newfoundland | 47.87 | -58.24 | 29 | NFL | 0 | 29 | 0 | TRUE | 1.00 | 0.235 | 0.228 |
| GBK | Garia Brook, Newfoundland | 47.75 | -58.53 | 29 | NFL | 0 | 29 | 0 | TRUE | 1.00 | 0.231 | 0.226 |
| FRM | Farmer's Brook, Newfoundland | 47.67 | -58.48 | 30 | NFL | 0 | 30 | 0 | TRUE | 1.00 | 0.253 | 0.249 |
| TYB | Humber River, Newfoundland | 49.55 | -57.10 | 29 | NFL | 0 | 28 | 1 | TRUE | 1.00 | 0.239 | 0.235 |
| LRBB | Lomond River Bond Bay Pond, Newfoundland | 49.36 | -57.65 | 23 | MAR | 0 | 0 | 23 | TRUE | 1.00 | 0.258 | 0.251 |
| CUS | Cuslett Brook, Newfoundland | 46.96 | -54.16 | 87 | NFL | 0 | 87 | 0 | FALSE | 0.31 | 0.227 | 0.232 |
| PHR | Pipers Hole River, Newfoundland | 47.93 | -54.27 | 86 | NFL | 0 | 86 | 0 | FALSE | 0.01 | 0.226 | 0.224 |
| BLA | Black River, Newfoundland | 47.89 | -54.17 | 82 | NFL | 0 | 80 | 2 | FALSE | 0.15 | 0.220 | 0.219 |
| LREB | Lomond River, Newfoundland East Branch, Newfoundland | 49.40 | -57.61 | 23 | MAR | 0 | 2 | 21 | TRUE | 1.00 | 0.246 | 0.240 |
| RHW | Red Harbour, Newfoundland River West | 47.30 | -55.02 | 75 | NFL | 0 | 75 | 0 | FALSE | 0.01 | 0.226 | 0.224 |
| WBSB | Western Brook Stag Brook, Newfoundland | 49.75 | -57.87 | 28 | MAR | 0 | 5 | 23 | TRUE | 1.00 | 0.240 | 0.234 |
| FLB | Flat Bay Brook, Newfoundland | 48.41 | -58.58 | 24 | MAR | 0 | 0 | 24 | TRUE | 1.00 | 0.241 | 0.238 |
| SHA | Sandy Harbour River, Newfoundland | 47.71 | -54.36 | 64 | NFL | 0 | 64 | 0 | FALSE | 0.02 | 0.229 | 0.228 |
| a_Mars | Ã Mars, Quebec | 48.34 | -70.87 | 26 | MAR | 0 | 0 | 26 | FALSE | 0.00 | 0.241 | 0.239 |
| COD | Grand Codroy, Newfoundland | 47.84 | -59.20 | 57 | MAR | 0 | 0 | 57 | TRUE | 0.98 | 0.232 | 0.231 |
| LWN | Lawn River, Newfoundland | 46.95 | -55.54 | 80 | NFL | 0 | 80 | 0 | FALSE | 0.00 | 0.227 | 0.228 |
| CBC | Come By Chance River, Newfoundland | 47.97 | -53.96 | 61 | NFL | 0 | 61 | 0 | FALSE | 0.02 | 0.231 | 0.234 |
| NMB | Northwest Brook, Newfoundland | 47.17 | -55.32 | 87 | NFL | 0 | 87 | 0 | FALSE | 0.24 | 0.225 | 0.229 |
| PBR | Piercey's Brook, Newfoundland | 46.88 | -55.86 | 83 | NFL | 0 | 83 | 0 | FALSE | 0.01 | 0.236 | 0.235 |
| GBW | Great Barasway Brook, Newfoundland | 47.12 | -54.06 | 89 | NFL | 0 | 89 | 0 | FALSE | 0.02 | 0.247 | 0.244 |
| TNR | Terra Nova River, Newfoundland | 48.67 | -54.00 | 29 | NFL | 0 | 29 | 0 | FALSE | 0.00 | 0.230 | 0.234 |
| KED | Kedgwick, New Brunswick | 47.91 | -67.91 | 15 | MAR | 0 | 0 | 15 | FALSE | 0.00 | 0.261 | 0.254 |
| JU | Jupiter, Quebec | 49.47 | -63.58 | 28 | MAR | 0 | 0 | 28 | FALSE | 0.00 | 0.238 | 0.238 |
| NRP | Northeast Placentia River, Newfoundland | 47.29 | -53.80 | 28 | NFL | 0 | 28 | 0 | FALSE | 0.04 | 0.247 | 0.241 |
| TRF | Trout River Feeder, Newfoundland | 49.65 | -57.76 | 29 | MAR | 0 | 0 | 29 | FALSE | 0.07 | 0.233 | 0.227 |
| BDN | Baydu Nord, Newfoundland | 47.73 | -55.44 | 19 | NFL | 0 | 19 | 0 | FALSE | 0.05 | 0.251 | 0.244 |
| UPS | Upsalquitch, New Brunswick | 47.57 | -66.54 | 28 | MAR | 0 | 0 | 28 | FALSE | 0.00 | 0.233 | 0.230 |
| BDL | Bayde L'Eau River, Newfoundland | 47.51 | -54.73 | 91 | NFL | 0 | 91 | 0 | FALSE | 0.00 | 0.225 | 0.230 |
| PAT | Patapedia, Quebec | 47.86 | -67.39 | 24 | MAR | 0 | 0 | 24 | FALSE | 0.00 | 0.241 | 0.237 |
| MAT | Matapedia, Quebec | 48.18 | -67.14 | 15 | MAR | 0 | 0 | 15 | FALSE | 0.00 | 0.261 | 0.252 |
| BRA | Branch River, Newfoundland | 46.89 | -53.97 | 67 | NFL | 0 | 67 | 0 | FALSE | 0.00 | 0.262 | 0.259 |
| TRB | Tailrace Brook, Newfoundland | 48.01 | -55.79 | 25 | NFL | 0 | 19 | 6 | TRUE | 0.80 | 0.244 | 0.233 |
| MAD | Madeleine, Quebec | 49.23 | -65.32 | 28 | MAR | 0 | 0 | 28 | FALSE | 0.00 | 0.237 | 0.234 |
| TREB | Trout River Eastern Brook, Newfoundland | 49.37 | -57.96 | 18 | MAR | 0 | 0 | 18 | FALSE | 0.17 | 0.281 | 0.262 |
| NPR | Northeast Placentia River, Newfoundland | 47.29 | -53.80 | 81 | NFL | 0 | 81 | 0 | FALSE | 0.07 | 0.238 | 0.237 |
| MOB | Mobile River, Newfoundland | 47.25 | -52.84 | 29 | NFL | 0 | 29 | 0 | FALSE | 0.00 | 0.252 | 0.239 |
| NEB19_YOY | Northeast Brook, Newfoundland | 47.73 | -55.36 | 28 | NFL | 0 | 28 | 0 | FALSE | 0.25 | 0.263 | 0.248 |
| CNR | Conne, Newfoundland | 47.91 | -55.70 | 202 | NFL | 0 | 202 | 0 | FALSE | 0.02 | 0.231 | 0.232 |
| TRWB | Trout River Western Brook, Newfoundland | 49.65 | -57.76 | 13 | MAR | 0 | 0 | 13 | FALSE | 0.23 | 0.269 | 0.256 |
| SHI | Ship Harbour Brook, Newfoundland | 47.35 | -53.87 | 82 | NFL | 0 | 82 | 0 | FALSE | 0.04 | 0.241 | 0.239 |
| NON | Nonsuch River, Newfoundland | 47.45 | -54.64 | 91 | NFL | 0 | 91 | 0 | FALSE | 0.00 | 0.236 | 0.233 |
| WCB | Western Cove Brook, Newfoundland | 47.51 | -53.68 | 17 | NFL | 0 | 17 | 0 | FALSE | 0.00 | 0.281 | 0.263 |
| GRB | Great Rattling Brook-Exploits, Newfoundland | 49.62 | -56.17 | 26 | NFL | 0 | 26 | 0 | FALSE | 0.00 | 0.242 | 0.238 |
| SHP | Sheepscot River, Maine USA | 43.91 | -69.67 | 20 | MAR | 0 | 0 | 20 | FALSE | 0.00 | 0.247 | 0.238 |
| NGR | Narraguagus, Maine USA | 44.52 | -67.86 | 21 | MAR | 0 | 0 | 21 | FALSE | 0.00 | 0.245 | 0.237 |
| LHR | Long Harbour, Newfoundland | 47.82 | -54.94 | 16 | NFL | 0 | 16 | 0 | FALSE | 0.00 | 0.270 | 0.260 |
| CMP | Campbellton, Newfoundland | 49.28 | -54.93 | 25 | NFL | 0 | 25 | 0 | FALSE | 0.00 | 0.246 | 0.241 |
| LBB | Little , Newfoundland Barasway Brook | 47.18 | -54.03 | 15 | NFL | 0 | 15 | 0 | FALSE | 0.00 | 0.298 | 0.277 |
| GBB1 | Grand Bank Brook, Newfoundland | 47.09 | -55.76 | 28 | NFL | 0 | 28 | 0 | FALSE | 0.04 | 0.261 | 0.244 |
| RNR | Renews River, Newfoundland | 46.94 | -52.96 | 30 | NFL | 0 | 30 | 0 | FALSE | 0.03 | 0.242 | 0.236 |
| GAR | Garnish, Newfoundland | 47.23 | -55.35 | 22 | NFL | 0 | 22 | 0 | FALSE | 0.00 | 0.266 | 0.255 |
| RUS | Rushoon River, Newfoundland | 47.37 | -54.92 | 84 | NFL | 0 | 84 | 0 | FALSE | 0.01 | 0.235 | 0.235 |
| AVR | Avondale River, Newfoundland | 47.39 | -53.23 | 29 | NFL | 0 | 29 | 0 | FALSE | 0.00 | 0.266 | 0.258 |
| TOB | Tobique River, New Brunswick | 46.77 | -67.70 | 25 | MAR | 0 | 0 | 25 | FALSE | 0.00 | 0.227 | 0.222 |
| NSH | Nashwaak, New Brunswick | 45.96 | -66.62 | 45 | MAR | 0 | 0 | 45 | FALSE | 0.00 | 0.210 | 0.208 |
| FHB | Fair Haven Brook, Newfoundland | 47.54 | -53.89 | 103 | NFL | 0 | 103 | 0 | FALSE | 0.00 | 0.254 | 0.249 |
| TDS | Tides Brook, Newfoundland | 47.13 | -55.26 | 68 | NFL | 0 | 68 | 0 | FALSE | 0.00 | 0.258 | 0.256 |
| TBR | Taylor Bay Brook, Newfoundland | 46.88 | -55.71 | 79 | NFL | 0 | 79 | 0 | FALSE | 0.00 | 0.241 | 0.240 |
| SJO | Saint John River, New Brunswick | 45.25 | -66.04 | 26 | MAR | 0 | 0 | 26 | FALSE | 0.00 | 0.223 | 0.219 |
| NBT | North Brook Trepassy, Newfoundland | 46.74 | -53.36 | 25 | NFL | 0 | 25 | 0 | FALSE | 0.00 | 0.287 | 0.276 |
| SER | Serpentine River, Newfoundland | 48.94 | -58.51 | 26 | MAR | 0 | 0 | 26 | FALSE | 0.00 | 0.232 | 0.227 |
| LMS2020 | Lamaline River, Newfoundland | 46.88 | -55.78 | 28 | NFL | 0 | 28 | 0 | FALSE | 0.00 | 0.251 | 0.245 |
| BSR | Big Salmon, New Brunswick | 45.42 | -65.41 | 98 | MAR | 0 | 0 | 98 | FALSE | 0.00 | 0.214 | 0.213 |
| RHA | Red Harbour River East, Newfoundland | 47.33 | -54.99 | 91 | NFL | 0 | 91 | 0 | FALSE | 0.00 | 0.253 | 0.252 |
| CLR | Collinet River, Newfoundland | 47.22 | -53.55 | 21 | NFL | 0 | 21 | 0 | FALSE | 0.00 | 0.270 | 0.263 |
| MAG | Magaudavic River, New Brunswick | 45.12 | -66.85 | 16 | MAR | 0 | 0 | 16 | FALSE | 0.00 | 0.272 | 0.257 |
| BCB | Bear Cove Brook, Newfoundland | 47.67 | -59.30 | 29 | MAR | 0 | 0 | 29 | FALSE | 0.00 | 0.261 | 0.246 |
| SLR | Salmonier River, Newfoundland | 47.20 | -53.37 | 30 | NFL | 0 | 30 | 0 | FALSE | 0.00 | 0.265 | 0.254 |
| SPR | Southeast Placentia River, Newfoundland | 47.23 | -53.88 | 96 | NFL | 0 | 96 | 0 | FALSE | 0.00 | 0.247 | 0.244 |
| LCO | Little Codroy, Newfoundland | 47.83 | -59.15 | 27 | MAR | 0 | 0 | 27 | FALSE | 0.00 | 0.242 | 0.236 |
| USR | Upper Salmon River, New Brunswick | 45.61 | -64.96 | 15 | MAR | 0 | 0 | 15 | FALSE | 0.00 | 0.278 | 0.264 |
| MED | Medway River, Nova Scotia | 44.14 | -64.64 | 23 | MAR | 0 | 0 | 23 | FALSE | 0.00 | 0.249 | 0.240 |
| LAH | La Have, Nova Scotia | 44.37 | -64.50 | 47 | MAR | 0 | 0 | 47 | FALSE | 0.00 | 0.213 | 0.213 |
| GLD | Gold River, Nova Scotia | 44.55 | -64.33 | 26 | MAR | 0 | 0 | 26 | FALSE | 0.00 | 0.256 | 0.246 |
| LSR | Little Salmonier, Newfoundland | 47.07 | -55.18 | 17 | NFL | 0 | 17 | 0 | FALSE | 0.00 | 0.271 | 0.263 |
| MUN | Miramichi-Upper Northwest, New Brunswick | 47.17 | -65.94 | 24 | MAR | 0 | 0 | 24 | FALSE | 0.00 | 0.238 | 0.233 |
| GAK | Gaspereau River, Nova Scotia | 45.06 | -64.38 | 111 | MAR | 0 | 0 | 111 | FALSE | 0.00 | 0.245 | 0.242 |
| MSW | Miramichi-Upper Southwest | 46.55 | -66.04 | 23 | MAR | 0 | 0 | 23 | FALSE | 0.00 | 0.243 | 0.235 |
| WES | West River Sheet Harbour, Nova Scotia | 44.95 | -62.59 | 53 | MAR | 0 | 0 | 53 | FALSE | 0.00 | 0.253 | 0.245 |
| SMA | St. Mary’s River, Nova Scotia | 45.10 | -61.96 | 26 | MAR | 0 | 0 | 26 | FALSE | 0.00 | 0.229 | 0.227 |
| NRH | North River, Nova Scotia | 45.38 | -63.31 | 22 | MAR | 0 | 0 | 22 | FALSE | 0.00 | 0.253 | 0.247 |
| KOU | Kouchibouguac, New Brunswick | 46.74 | -65.20 | 31 | MAR | 0 | 0 | 31 | FALSE | 0.00 | 0.230 | 0.228 |
| MOR | Morells, Prince Edward Island | 46.30 | -62.71 | 18 | MAR | 0 | 0 | 18 | FALSE | 0.00 | 0.272 | 0.260 |
| CHT | Cheticamp River, Nova Scotia | 46.64 | -60.95 | 12 | MAR | 0 | 0 | 12 | FALSE | 0.00 | 0.272 | 0.263 |
| NET | Northeast, Prince Edward Island | 46.38 | -62.57 | 24 | MAR | 0 | 0 | 24 | FALSE | 0.00 | 0.256 | 0.253 |
| RIC | Richibucto, New Brunswick | 46.36 | -65.15 | 31 | MAR | 0 | 0 | 31 | FALSE | 0.00 | 0.235 | 0.232 |
| SCP | South Central, Prince Edward Island | 46.28 | -63.49 | 14 | MAR | 0 | 0 | 14 | FALSE | 0.00 | 0.295 | 0.273 |
| NASP | North Aspy, Nova Scotia | 46.83 | -60.61 | 29 | MAR | 0 | 0 | 29 | FALSE | 0.00 | 0.244 | 0.236 |
| RPH | River Philip, Nova Scotia | 45.59 | -63.82 | 17 | MAR | 0 | 0 | 17 | FALSE | 0.00 | 0.255 | 0.247 |
| CLY | Clyburne, Nova Scotia | 46.66 | -60.41 | 28 | MAR | 0 | 0 | 28 | FALSE | 0.00 | 0.243 | 0.239 |
| MNE | Northeast Margaree, Nova Scotia | 46.47 | -60.92 | 12 | MAR | 0 | 0 | 12 | FALSE | 0.00 | 0.275 | 0.264 |
| BAD | Baddeck, Nova Scotia | 46.10 | -60.84 | 28 | MAR | 0 | 0 | 28 | FALSE | 0.00 | 0.243 | 0.238 |
| NWP | Northwest Complex Prince Edward Island | 46.63 | -64.04 | 17 | MAR | 0 | 0 | 17 | FALSE | 0.00 | 0.273 | 0.265 |
| NEP | Northeast Complex Prince Edward Island | 46.45 | -62.21 | 27 | MAR | 0 | 0 | 27 | FALSE | 0.00 | 0.278 | 0.262 |
| PIE | East River Pictou, Nova Scotia | 45.54 | -62.88 | 23 | MAR | 0 | 0 | 23 | FALSE | 0.00 | 0.245 | 0.241 |
| ING | Ingonish, Nova Scotia | 46.62 | -60.45 | 29 | MAR | 0 | 0 | 29 | FALSE | 0.00 | 0.243 | 0.237 |
| STW | Stewiacke, Nova Scotia | 45.14 | -63.38 | 102 | MAR | 0 | 0 | 102 | FALSE | 0.00 | 0.254 | 0.252 |
| MRS | Southwest Margaree, Nova Scotia | 46.24 | -61.12 | 14 | MAR | 0 | 0 | 14 | FALSE | 0.00 | 0.288 | 0.265 |
| JGC | Graham River, Nova Scotia | 45.86 | -61.49 | 11 | MAR | 0 | 0 | 11 | FALSE | 0.00 | 0.285 | 0.274 |
| MAB | Mabou River, Nova Scotia | 46.04 | -61.31 | 27 | MAR | 0 | 0 | 27 | FALSE | 0.00 | 0.251 | 0.245 |
| INH | Inhabitants River, Nova Scotia | 45.60 | -61.23 | 27 | MAR | 0 | 0 | 27 | FALSE | 0.00 | 0.261 | 0.244 |

Table S2. Summary of genomic windows with evidence of high linkage disequilibrium.

| chromosome | Window start | N loci | N comparisons | Median r^2^ | cluster |
| --- | --- | --- | --- | --- | --- |
| 2 | 45000001 | 4 | 7 | 0.717354 | Newfoundland |
| 3 | 14000001 | 7 | 13 | 0.908362 | full |
| 8 | 9000001 | 4 | 5 | 1 | Labrador |
| 9 | 1000001 | 8 | 17 | 0.737871 | Maritimes |
| 9 | 41000001 | 6 | 17 | 1 | full |
| 9 | 46000001 | 5 | 7 | 0.740569 | Maritimes |
| 10 | 28000001 | 5 | 11 | 1 | full |
| 10 | 1.15E+08 | 4 | 5 | 0.680423 | Newfoundland |
| 11 | 92000001 | 4 | 6 | 0.688074 | Newfoundland |
| 12 | 21000001 | 8 | 16 | 0.842818 | Labrador |
| 15 | 49000001 | 6 | 11 | 1 | full |
| 18 | 9000001 | 8 | 24 | 1 | full |
| 19 | 48000001 | 7 | 10 | 0.9771 | full |
| 19 | 51000001 | 10 | 41 | 1 | full |
| 19 | 52000001 | 4 | 12 | 0.872956 | full |
| 19 | 61000001 | 4 | 10 | 0.663581 | Newfoundland |
| 20 | 41000001 | 3 | 5 | 0.810302 | Maritimes |
| 23 | 1 | 8 | 27 | 0.944812 | full |
| 23 | 1 | 8 | 30 | 0.944812 | Labrador |
| 23 | 1000001 | 7 | 32 | 0.885357 | full |
| 23 | 1000001 | 7 | 40 | 0.882061 | Labrador |
| 24 | 20000001 | 4 | 5 | 0.722299 | Newfoundland |
| 24 | 44000001 | 4 | 7 | 1 | full |
| 26 | 1000001 | 9 | 16 | 0.763754 | Maritimes |

Table S3. Heterozygosity summary statistics for the three populations and complete set of individuals.

| Population | Number of samples | Number of locations | Mean observed heterozygosity | Mean expected heterozygosity |
| --- | --- | --- | --- | --- |
| Complete data set | 5451 | 149 | 0.184 | 0.201 |
| Labrador | 431 | 20 | 0.212 | 0.228 |
| Newfoundland | 3244 | 71 | 0.201 | 0.213 |
| Maritimes | 1776 | 69 | 0.189 | 0.202 |

Table S4. Genes that were identified in two of the following analyses: i) contained one or more SNPs identified as significant in the PCA analysis, ii) that contained one or more SNPs within identified as an Fst outlier in one of the pairwise comparisons, ii) that fell within a window of significant linkage disequilibrium.

| Gene Name | Evidence | Atlantic salmon chromosome number | gene bp start | gene bp end | Gene ID |
| --- | --- | --- | --- | --- | --- |
| GDNF family receptor alpha 4a | fst and LD | 1 | 1586801 | 1713084 | 106571988 |
| cysteine-rich motor neuron 1 protein-like | fst and PCA | 1 | 25647662 | 25696085 | 106586563 |
| formin-1 | fst and PCA | 1 | 25764293 | 25861942 | 106586878 |
| ryanodine receptor 3 | fst and PCA | 1 | 25872092 | 26041001 | 106587081 |
| regulator of G-protein signaling 6 | fst and PCA | 1 | 26494902 | 26569238 | 106588600 |
| ligand-dependent corepressor | fst and PCA | 1 | 77893068 | 77935594 | 106612867 |
| astrotactin-2-like | fst and LD | 1 | 140201272 | 1.41E+08 | 106565912 |
| myocyte-specific enhancer factor 2C-like | fst and PCA | 1 | 150969950 | 1.51E+08 | 106568324 |
| glutamate receptor ionotropic, kainate 2 | fst and PCA | 2 | 27135446 | 27405776 | 106578944 |
| protein cordon-bleu-like | fst and PCA | 2 | 31229859 | 31308089 | 106579857 |
| growth factor receptor-bound protein 10-like | fst and PCA | 2 | 31316626 | 31407281 | 106579904 |
| heparan sulfate glucosamine 3-O-sulfotransferase 4-like | fst and PCA | 3 | 63033581 | 63130221 | 106601107 |
| neurofibromin 1b | fst and PCA | 4 | 47300222 | 47373907 | 106603288 |
| RNA-binding protein Musashi homolog 2 | fst and PCA | 4 | 47403212 | 47706787 | 106603292 |
| delta-sarcoglycan-like | fst and LD | 4 | 58477430 | 58589395 | 106603489 |
| rho GTPase-activating protein 7-like | fst and LD | 4 | 58658555 | 58746828 | 106603488 |
| protein KIBRA-like | fst and LD | 4 | 58788039 | 58832923 | 123742473 |
| G protein-coupled receptor kinase 6-like | fst and LD | 4 | 58850547 | 58893636 | 106603485 |
| proline-rich protein 7 | fst and LD | 4 | 58914157 | 58927820 | 106603484 |
| rho GTPase-activating protein 26 | fst and LD | 4 | 76886801 | 77058905 | 106603859 |
| ras-related protein Rab-13 | fst and PCA | 5 | 75081702 | 75093760 | 123743170 |
| glucagon receptor-like | fst and LD | 6 | 8542489 | 8836467 | 106594483 |
| DENN domain-containing protein 5B | fst and PCA | 7 | 41962468 | 42040924 | 106609315 |
| adenosine deaminase 2b | fst and PCA | 7 | 46032538 | 46037525 | 106609445 |
| F-box/WD repeat-containing protein 7 | fst and PCA | 8 | 12467038 | 12671648 | 106610174 |
| WD repeat domain 25 | fst and PCA | 9 | 6554577 | 6597024 | 106610588 |
| disintegrin and metalloproteinase domain-containing protein 17 | fst and PCA | 9 | 11284332 | 11306796 | 106610723 |
| charged multivesicular body protein 3 | fst and PCA | 9 | 23339965 | 23345673 | 106610994 |
| metabotropic glutamate receptor 8 | fst and PCA | 10 | 77099384 | 77362422 | 106561032 |
| WD repeat domain 43 | fst and PCA | 11 | 1027815 | 1077742 | 106561875 |
| neural-cadherin | fst and PCA | 11 | 13716361 | 13972076 | 106562040 |
| calmodulin-binding transcription activator 1-like | fst and PCA | 13 | 34675147 | 35161033 | 106567259 |
| uncharacterized LOC106567691 | fst and PCA | 13 | 72819685 | 72895900 | 106567691 |
| AF4/FMR2 family member 1-like | fst and PCA | 13 | 106136726 | 1.06E+08 | 106568507 |
| NACHT, LRR and PYD domains-containing protein 1 homolog | fst and LD | 14 | 43071019 | 43093935 | 106569585 |
| natural killer cell receptor 2B4-like | PCA and LD | 14 | 43212879 | 43298705 | 106592761 |
| uncharacterized LOC123726710 | PCA and LD | 14 | 43246491 | 43262963 | 123726710 |
| rho guanine nucleotide exchange factor 18 | fst and LD | 14 | 43843325 | 43881286 | 106569603 |
| ankyrin repeat and SAM domain-containing protein 1A | fst and PCA | 15 | 83915689 | 84034309 | 106572349 |
| glypican-6-like | fst and LD | 17 | 45112031 | 45472099 | 106575058 |
| protein eyes shut homolog | fst and PCA | 18 | 33751045 | 33973429 | 106577282 |
| cadherin-12 | fst and LD | 19 | 21099165 | 21349524 | 106578565 |
| cpn1 carboxypeptidase N, polypeptide 1 | fst and LD | 19 | 66828633 | 67011882 | 100196707 |
| calcitonin gene-related peptide type 1 receptor | fst and PCA | 21 | 21556592 | 21615393 | 106581932 |
| calcium/calmodulin-dependent 3',5'-cyclic nucleotide phosphodiesterase 1A | fst and LD | 21 | 34913509 | 35014822 | 106582264 |
| FERM domain-containing protein 4B-like | fst and PCA | 22 | 15082728 | 15146984 | 106582859 |
| potassium voltage-gated channel subfamily C member 1 | fst and PCA | 23 | 3373044 | 3499951 | 106583948 |
| insulin receptor-like | fst and PCA | 23 | 3543987 | 3696146 | 106583950 |
| inactive tyrosine-protein kinase transmembrane receptor ROR1 | fst and PCA | 23 | 9307191 | 9500357 | 106584030 |
| peripheral plasma membrane protein CASK | fst and PCA | 25 | 41006591 | 41237760 | 106586786 |
| tent2 | fst and PCA | 26 | 4174984 | 4216958 | 106587048 |
| homer scaffold protein 1b | fst and PCA | 26 | 4250234 | 4430123 | 106587049 |
| protein kinase C-binding protein NELL1 | fst and PCA | 26 | 5916821 | 6410687 | 106587057 |
| thyroid hormone receptor beta | fst and PCA | 27 | 20294180 | 20432700 | 106588669 |
| myocardin | fst and PCA | 28 | 12183786 | 12331195 | 106589488 |
| growth arrest-specific 7b | fst and PCA | 28 | 12387833 | 12440508 | 106589492 |
| glucagon-like peptide 2 receptor | fst and PCA | 28 | 12516465 | 12535766 | 106589495 |

Table S5. Pathways of interest identified through Gene Ontology analysis.

| Enrichment FDR | nGenes | Pathway Genes | Fold Enrichment | Pathways |
| --- | --- | --- | --- | --- |
| 4.00E-02 | 2 | 31 | 70.9 | Cellular calcium ion homeostasis |
| 4.30E-02 | 2 | 36 | 61 | Cellular divalent inorganic cation homeostasis |
| 4.40E-02 | 2 | 40 | 54.9 | Calcium ion homeostasis |
| 4.80E-02 | 2 | 45 | 48.8 | Divalent inorganic cation homeostasis |
| 1.60E-02 | 4 | 239 | 18.4 | Transmembrane receptor protein tyrosine kinase signaling pathway |
| 4.00E-02 | 4 | 345 | 12.7 | Enzyme linked receptor protein signaling pathway |
| 1.60E-02 | 7 | 1153 | 6.7 | Cell surface receptor signaling pathway |
